# T Cell Receptor Beta Germline Variability is Revealed by Inference From Repertoire Data

**DOI:** 10.1101/2021.05.17.444409

**Authors:** Aviv Omer, Ayelet Peres, Oscar L Rodriguez, Corey T Watson, William Lees, Pazit Polak, Andrew M Collins, Gur Yaari

**Author notes:** Equal contributor (Aviv Omer, Ayelet Peres).

## Abstract

**Background:** T and B cell receptor (TCR, BCR) repertoires constitute the foundation of adaptive immunity. Adaptive immune receptor repertoire sequencing (AIRR-seq) is a common approach to study immune system dynamics. Understanding the genetic factors influencing the composition and dynamics of these repertoires is of major scientific and clinical importance. The chromosomal loci encoding for the variable regions of TCRs and BCRs are challenging to decipher due to repetitive elements and undocumented structural variants.

**Methods:** To confront this challenge, AIRR-seq-based methods have recently been developed for B cells, enabling genotype and haplotype inference and discovery of undocumented alleles. However, this approach relies on complete coverage of the receptors’ variable regions, whereas most T cell studies sequence a small fraction of that region. Here, we adapted a B cell pipeline for undocumented alleles, genotype, and haplotype inference for full and partial TCR sequences. The pipeline also deals with gene assignment ambiguities, which is especially important in the analysis of data-sets of partial sequences.

**Results:** We identified 39 undocumented polymorphisms in T cell receptor Beta V (TRBV) and 31 undocumented 5’ UTR sequences. A subset of these inferences was also observed using independent genomic approaches. We found that a single nucleotide polymorphism differentiating between the two documented T cell receptor Beta D2 (TRBD2) alleles is strongly associated with dramatic changes in the expressed repertoire.

**Conclusions:** We reveal a rich picture of germline variability, and demonstrate how a single nucleotide polymorphism dramatically affects the composition of the whole repertoire. Our findings provide a basis for annotation of TCR repertoires for future basic and clinical studies.

## Background

The immune system’s success in fighting countless evolving pathogens depends on a dynamic and diverse set of B and T cell receptors. Due to the longevity of immunological memory, high throughput sequencing of adaptive immune receptor repertoires (AIRR-seq) provides detailed insights into the past and present encounters of the human immune system [1]. It can teach us about fundamental immune processes and reveal dysregulation, with broad implications for biomedicine. B and T cell receptors are assembled within B and T cells, respectively, during differentiation from hematopoietic stem cells, by a complex process involving somatic recombination of a large number of germline-encoded Variable (V), Diversity (D), and Joining (J) gene segments, along with junctional diversity in the form of addition and subtraction of nucleotides at the boundaries where these segments are joined together [2]. This V(D)J recombination process creates a diverse repertoire of receptors that together with the innate immune system form the first line of defense against pathogens.

Genetic factors are expected to influence the structure and functionality of AIRRs [3, 4]. However, understanding these genetic effects is confounded by lack of knowledge about the population genetics of the TCR and BCR encoding genomic loci, and the special challenges involved in describing the germline gene set of any individual [5]. The lack of knowledge about these loci is due to difficulties in reliably mapping repetitive elements and undocumented structural variations with short read sequencing. Many TCR genes were reported in the decade after their first discovery [6]. Complete sequences of the TCR encoding loci were reported in 1985 [7–9], and this led to the development of the TCR encoding gene nomenclature by the ImMunoGeneTics (IMGT) group [10]. Since then, very few allelic variants have been entered into the IMGT reference directories of germline genes. In fact no new allelic variants of the TCR variable region genes have been named this century despite published studies suggesting that the IMGT TCR reference directory may be far from complete [11–13].

Until recently, there has been a similar lack of attention paid to the documentation of BCR encoding loci, because the direct genomic sequencing of these loci is also very challenging [14, 15]. This changed with the development of a method for targeted long-read direct sequencing of the BCR heavy chain encoding locus [16], and with the realisation that BCR encoding undocumented germline alleles and genotypes can be reliably inferred from AIRR-seq data [17–20], as well as haplotypes [21, 22], and chromosomal deletions within the BCR encoding loci [23]. Even though TCR V(D)J gene rearrangements are generated by analogous mechanisms to BCR rearrangements, to date there is no published data about TCR germline allele inference and structural variation in the TCR encoding loci. Recently, there were attempts to extract BCR and TCR encoding allelic information from short read whole genome sequencing data [11, 24, 25], but these approaches were not validated with targeted sequencing or AIRR-seq, and therefore are subject to criticism regarding the reliability of the inferences [5, 26]. Hence, to study genomic variations in TCR encoding loci and their relations to the expressed repertoires, there is a need to adapt BCR inference tools to TCR data.

In T cells, due to lack of somatic mutations, most studies sequence only a small fraction of the variable region. AIRR-seq data can be generated by methods that differ in the length of their coverage of V(D)J sequences. 5’ RACE amplifies the whole V(D)J region from the 3’ end of the J region to the 5’ end of the mRNA molecule. BIOMED-2 primers [27] amplify partial VDJ sequences from the J gene to the framework-2 (FR2) of the V gene, while the Adaptive Biotechnologies [28] approach generates only 87 nucleotides from a fixed position within each T cell receptor Beta J (TRBJ) gene in FR4, and includes the complementary determining region 3 (CDR3) and a fraction of the TRBV gene from FR3. As there are TRBV alleles that are identical to other TRBV alleles for varying lengths from their 3’ ends, in some cases it is impossible to identify the gene and allele source of these partial V(D)J sequences, generating a serious gene assignment ambiguity problem. Thus, the development of a TRBV genotype inference method requires the detailed documentation of any gene pairs that can be impossible to distinguish for a given sequencing approach.

Here, we adapt a B cell inference pipeline to TRB AIRR-seq data (Fig. 1A), to work with these three types of sequencing approaches (Fig. 1B), and infer undocumented alleles, i.e., alleles that were previously not documented in IMGT, as well as single and double chromosome deletions, genotypes, and haplotypes. The pipeline was adapted to deal with the gene assignment ambiguity problem, which is especially important in the analysis of data-sets of partial sequences (Fig. 1C-D). This is an even greater problem in data-sets of partial sequences (Fig. 1C-D). We applied this pipeline to four of the largest AIRR-seq data-sets currently available and revealed a rich picture of germline variability, and a demonstration of how a single nucleotide polymorphism dramatically affects the composition of the whole repertoire. TCR Germline variability and its effects on the expressed repertoire may have important implications for TCR based immunotherapy and disease diagnosis, for example by enabling the discovery of TCR-related predispositions to diseases, or responses to different therapies.

**Figure 1.**
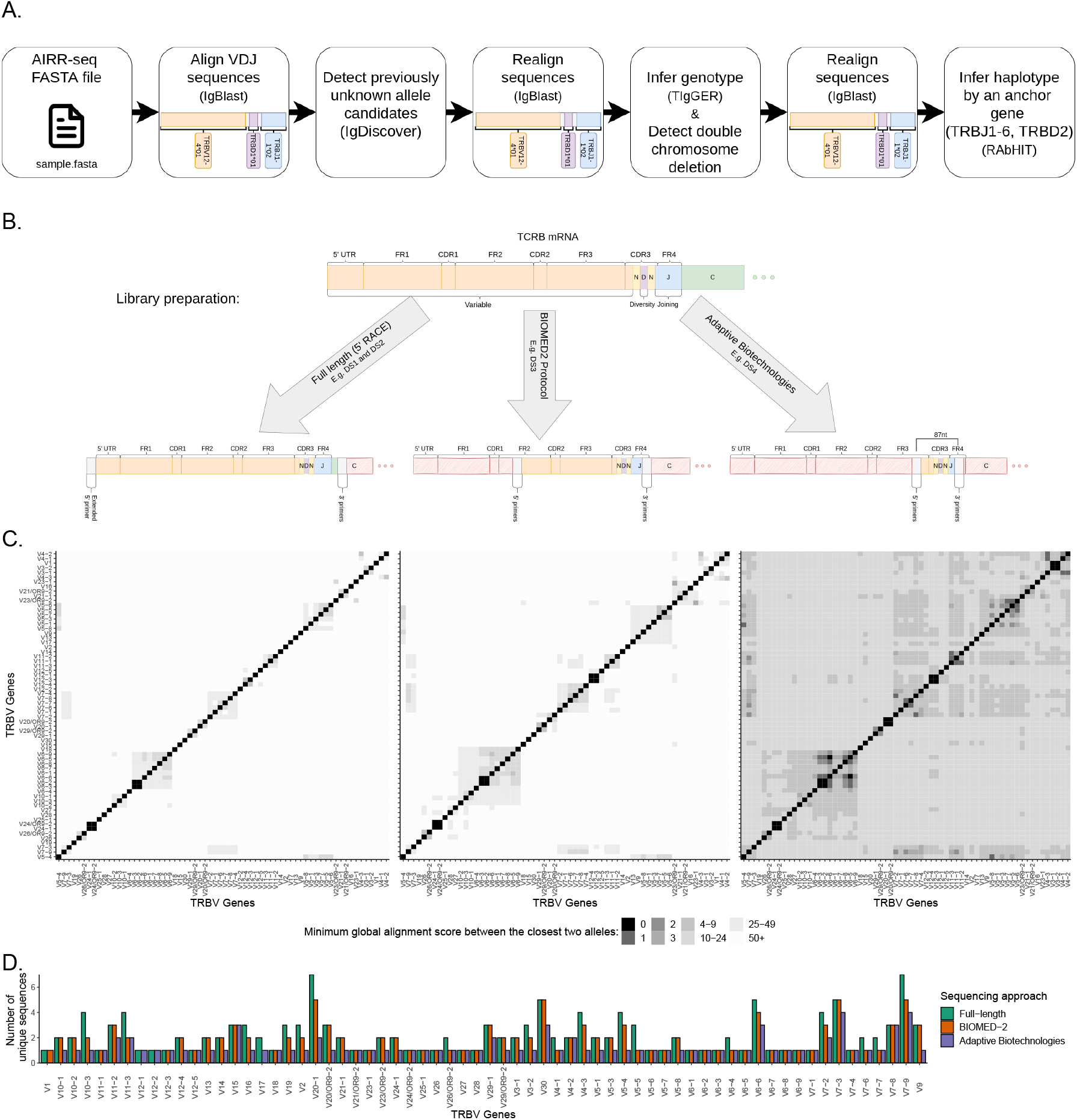
Overview of TRB genotyping and haplotyping workflow, TCR*β* sequencing technologies, and number of documented alleles. (A) An illustration of the steps for TRB genotyping and haplotyping. The input is an AIRR-seq fasta file after preprocessing, according to the sequencing protocol described in the methods (see section *Data*). (B) An illustration of the three TCR*β* sequencing approaches investigated in this study. (C) Heatmaps of distances between amplified TRBV gene sequences. The three panels correspond to the sequencing approaches shown above them. Colors correspond to the distances between the genes’ as measured by the minimum global alignment score (with a padded beginning and a penalty of one for each mismatch, insertion, or deletion) between the closest two alleles in the sequenced region. (D) Barplot of the unique sequence number per gene in the studied sequencing approaches.

## Methods

### Data

Four data-sets (DSs) were collected [29–34] (Table 1). Three were bulk sequenced, but each of them was generated using a different sequencing protocol, producing sequences of different lengths. The DSs also differ in sequencing depth, i.e., the number of unique sequences per repertoire (Additional file 1:Fig. S1). Another DS of TCRs amplified from single-cells was collected from three different sources [33–35], see Additional file 1:Table S1. The data-sets are described in Table. 1. In DS3 there were originally 313 individuals and 348 samples, out of which 108 samples were obtained from 94 individuals with hematological cancer. These 108 samples were dropped out of the analysis, given their likely bias towards substantial mutation and oligoclonality. DS2 and DS4 were downloaded after preprocessing, DS1 was preprocessed according to the preprocessing of Eliyahu et al. [29], and DS3 was preprocessed using pRESTO [36] according to the example workflow “Illumina MiSeq 2×250 BCR mRNA” as follows, (i) paired-ends were assembled, (ii) sequences with low quality (mean Phred quality scores lower than 20) were removed, (iii) the 3’ and 5’ end primers were cut, (iv) duplicate sequences were removed and collapsed.

**Table 1.**
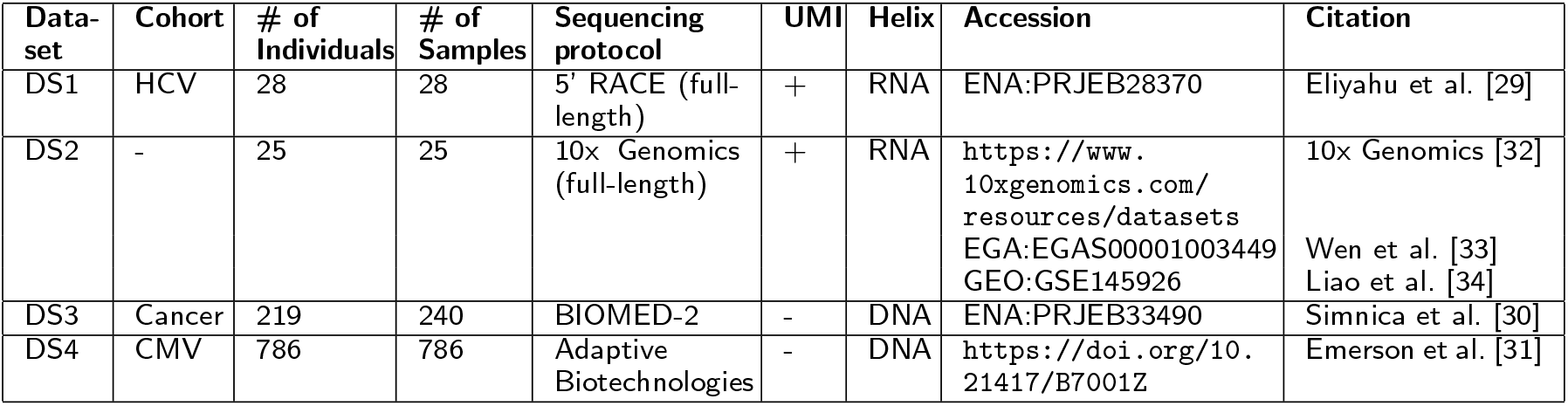
Data-sets (DS) analyzed in this study.

### Merging indistinguishable genes

Sequences of two full-length TRBV genes, TRBV6-2*01 and TRBV6-3*01, are indistinguishable (Fig. 1C). We therefore refer to them here as TRBV6-2*01/TRBV6-3*01. TRBV sequences amplified using the BIOMED-2 primers are partial, yet it is still possible to differentiate most of the genes. Only TRBV6-2 and TRBV6-3, as well as TRBV12-3 and TRBV12-4 could not be differentiated (Fig. 1C). Those indistinguishable sequences are referred to here as TRBV6-2/TRBV6-3 and TRBV12-3/TRBV12-4. The Adaptive Biotechnologies sequencing protocol generates very short partial TRBV gene sequences, yet it is still possible to identify most of them. Only the gene pairs TRBV6-2/TRBV6-3, TRBV12-3/TRBV12-4, TRBV3-1/TRBV3-2 and TRBV6-5/TRBV6-6 were indistinguishable. (Fig. 1C).

Adaptive Biotechnologies supplies 87 nucleotides from a fixed position within each TRBJ gene [28]. As a result, the given coverage of the TRBV segment is not constant, because TRBJ genes and junction regions have different lengths. Therefore, the distribution of the first position that the sequences covered of the TRBV reference was investigated. 96% of the sequences cover the first position following the TRBV gene primers. The BIOMED-2 protocol did not include primers for TRBV12-2. Thus, we were unable to explore the usage or genetic variation of this gene in DS3.

### Allele pattern collapsing

Although there are few ambiguities in the identification of partial TRBV genes, the unambiguous identification of partial allelic variants is more problematic. Many SNPs that distinguish between alleles are located outside the regions that are generated using BIOMED-2 or Adaptive Biotechnologies primers. Thus, all alleles were collapsed into partial allelic variation groups, the sequence of each partial allelic variation group was determined to be identical to the longest allele sequence reference (out of the identical partial alleles’ references). The allele patterns were named here using the following structure: [gene name]*[protocol primers][0-9][0-9]. The BIOMED-2 partial allelic variants were symbolized by bp, and the Adaptive Biotechnologies partial allelic variants were symbolized by ap. For example, the partial sequence of the allele TRBV5-6*01 was collapsed into the partial allelic variation groups TRBV5-6*bp01 and TRBV5-6*ap01 (see Additional file 1:Table S2 and Additional file 1:Table S3). The primers of the BIOMED-2 protocol were taken from van Dongen et al. [27], and the primers of Adaptive Biotechnologies were taken from Robins et al. [28].

### Genotype and undocumented allele inference

IgDiscover [17] was used for detection of undocumented allele candidates, and TIgGER for genotype inferences. Sequences were first aligned with IgBlast and processed using the IgDiscover igblast function and the Change-O MakeDb function [37]. A correction to the inferred CDR3 sequences in the IgDiscover output was done by replacing the sequences with their counterparts in the MakeDb output. TRBV allele candidates were inferred using IgDicover’s ‘‘discover” function. Undocumented allele candidates were filtered based on several rules. First, suspected SNPs were counted only between the boundaries: 5’ position of N+5 where N is the nucleotide position in which the primer ends or the sequence starts (the larger between the two), and 3’ position 316 by IMGT numbering. Second, candidate undocumented alleles were filtered out if they were not an exact match to at least 5% of the gene alignments. Third, such candidates had to have a sufficient rearrangement diversity: at least two different CDR3 lengths and two TRBJ genes. Fourth, for noisy data-sets in which chimeras were observed, candidates that could result from chimerism were filtered out. A candidate was suspected as having the potential to result from chimerism if two alleles from separate genes could generate an exact matched sequence in the range between nucleotides N+5 and 316.

Undocumented allele candidates were then combined with the IMGT TRBV Reference Directory to create individual-specific Reference Directories. Sequence sets were then re-aligned against the new directories, and genotypes were constructed with TIgGER using a Bayesian approach [19]. Genotyping was limited to sequences with a single assignment (only one best match), and with up to one mismatch in the TRBV segment. For the construction of the TRBD genotype, sequences with mismatches in the TRBD segment or with identifiable TRBD sequences shorter than nine nucleotides were filtered out. TIgGER’s level of confidence was calculated using a Bayes factor (K) from the posterior probability for each model. The larger the K, the greater the certainty in the genotype inference. lk that is used throughout the manuscript indicates the log of K.

For the undocumented alleles two additional filters after the genotype inferences were added. First, undocumented alleles were filtered out if there was more than one SNP within a stretch of four adjacent nucleotides [38]. Second, to account for potential sequencing errors, we investigated the modality of the undocumented alleles’ distribution in the population. The alleles that did not follow the expected usage distribution of bi or tri modal were discarded.

### Validating undocumented alleles/variants in long-read assemblies

35 diploid (70 haplotypes) HiFi long-read assemblies from 32 unrelated individuals and three offspring were downloaded from Ebert et al. [39] and aligned to GRCh38 [40] using BLASR with default parameters [41]. Gene sequences (5’ UTR, leader-1, leader-2 and exons) from the assemblies were extracted based on the alignment. Undocumented alleles and variants were determined to be present in the assemblies only if they exactly matched the extracted gene sequence.

### Determining the borders between TRBD2 genotype groups

We examined the fraction of TRBD2*01 assignments amongst all sequences unambiguously assigned to TRBD2 in DS1. The observed frequencies defined three distinct groups (Additional file 1:Fig. S2A). In the first group (~ 0 – 0.2), TRBD2*01 assignments peaked around 0.125 of the TRBD2 annotations. The second group (~ 0.2 – 0.8) peaked around 0.5, and the third group (~ 0.8 – 1) peaked around 0.95. According to this distribution, individuals in the first group are homozygous for TRBD2*02, and all alignments to TRBD2*01 are in error. The second group corresponds to individuals who are heterozygous for TRBD2, and individuals in the third group are homozygous for TRBD2*01. The resulting genotype frequency distribution of TRBD2 were in Hardy-Weinberg equilibrium.

To better define the thresholds differentiating the three groups, we turned to the larger data-sets. DS3 was too noisy for this purpose (Additional file 1:Fig. S2B), most likely due to the library preparation and sequencing protocol. In DS4, which contains 768 individuals, the TRBD2*01 distribution was tri-modal and very similar to that in DS1 (Additional file 1:Fig. S2C). The homozygous TRBD2*01 group is centered around a frequency of 0.96. The heterozygous group is centred around 0.45, and the homozygous TRBD2*02 group is centered around 0.12. The medians of the three groups are close to their means (Additional file 1:Table. S4), and as the size of the groups is large enough, we used the central limit theorem, and thus assume that the three data-sets are normally distributed.

Boundaries between the tri-modal peaks were defined as the equilibrium points between the peaks. The equilibrium point between two normal distributions is the point between the two averages of the groups, at which the probability of being in either one of the two groups is equal. For two distributions *X*_1_ ~ *N*_1_(*μ*_1_, *σ*_1_) and *X*_2_ ~ *N*_2_(*μ*_2_, *σ*_2_), the equilibrium point x is determined as follows:

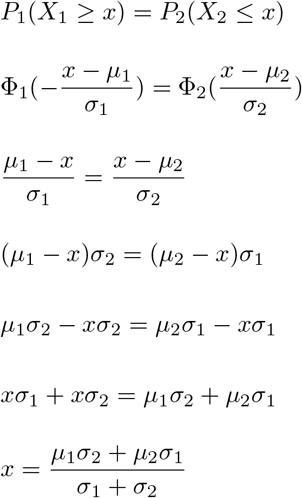

Applying the above to DS4 resulted in the following borders: the homozygous TRBD2*02 group was composed of 192 samples with a fraction of TRBD2*01 lower than 0.2066, the heterozygous group was composed of 404 samples with a fraction between 0.2066-0.8968, and the homozygous TRBD2*01 group was composed of 190 samples with a TRBD2*01 fraction above 0.8968. Thus, the D2 allele frequency was around 0.244 in homozygous individuals for TRBD2*01, 0.514 in heterozygous individuals, and 0.242 in homozygous individuals for TRBD2*02, consistent with the Hardy-Weinberg principle.

### Gene usage comparison

The differences in gene usage between groups were analyzed using a two-tailed Mann-Whitney test with the p value for significance adjusted by the Bonferroni correction to deal with the problem of multiple comparisons.

### Double chromosome deletion inference

The detection of double chromosome deletion was done using a published method [23], and adapted to TRB. Briefly, the method assesses whether a gene can be declared as deleted using a binomial test where the parameters for a given individual are: X is set to the number of sequences mapped to a given gene, N is the total number of sequences, and P is the lowest lowest relative frequency for the given gene from the non-deleted candidates. To determine a threshold for a given gene, a minimum cutoff is set and the closest frequency above the cutoff is set as (P). For TRB, the minimum cutoff of the average gene usage was lowered from 0.005 to 0.0005. A data frame table was used that contained the following columns: individual, gene, N, and total. N represents the number of unique sequences for each gene. The total column records the total number of the individual’s unique sequences. A binomial test for detecting chromosome deletions was then applied to the data frame table.

### Haplotype inference

RAbHIT was used as previously described [22], with TRBD2 and TRBJ1-6 anchors to infer TCR haplotypes. The epsilon error parameter was adjusted to deal with TRBD2 alignment errors, and was estimated with reference to the frequency distribution of TRBD2*01 alignments amongst all TRBD2-bearing sequences (see Additional file 1:Fig. S2) mentioned above. As ~12.5% of TRBD2*02 rearrangements mis-align to TRBD2*01, the epsilon, a parameter which defines the probability of the mis-assignment, was set at 0.125 if this allele dominated the TRBD2 alignments. Around 4% of the TRBD2*01 rearrangements mis-align to TRBD2*02, and epsilon was therefore set at 0.04 if TRBD2*01 dominated the alignments.

## Results

### Identification of undocumented alleles and upstream sequence variations within TRBV genotypes

To explore allelic variation in the TRBV locus, we inferred the sets of alleles carried for each expressed TRBV gene in many individuals, i.e., personal genotypes. For this, we took a multi step approach, using IgDiscover [17] and TIgGER [18], which yielded a set of candidate sequences that have not previously been documented in IMGT. Hereafter, we will refer to such sequences as “undocumented alleles”. To ensure our confidence in the inference of undocumented alleles, we took steps that reduce the influence of sequencing errors (see methods).

We applied the above approach to four data-sets, spanning different sequencing protocols and scales (see methods). From data-set 1 (DS1), which includes 28 individuals sequenced in full length, we inferred 18 undocumented alleles (Fig. 2A). Four of the 18 alleles were only seen at very low levels - less than 20% of all identified alleles of those genes, and showed a uni-modal distribution. This did not follow our expected usage distribution (these alleles are marked red in Fig. 2A). These candidate “alleles” therefore potentially result from sequencing errors. Two more undocumented alleles were considered erroneous due to the presence of more than one SNP in short nucleotide stretches. After discarding these six allele candidates, we were left with 12 undocumented alleles. Nine of the alleles were observed in more than a single individual’s genotype, which increases our confidence in the inferences.

**Figure 2.**
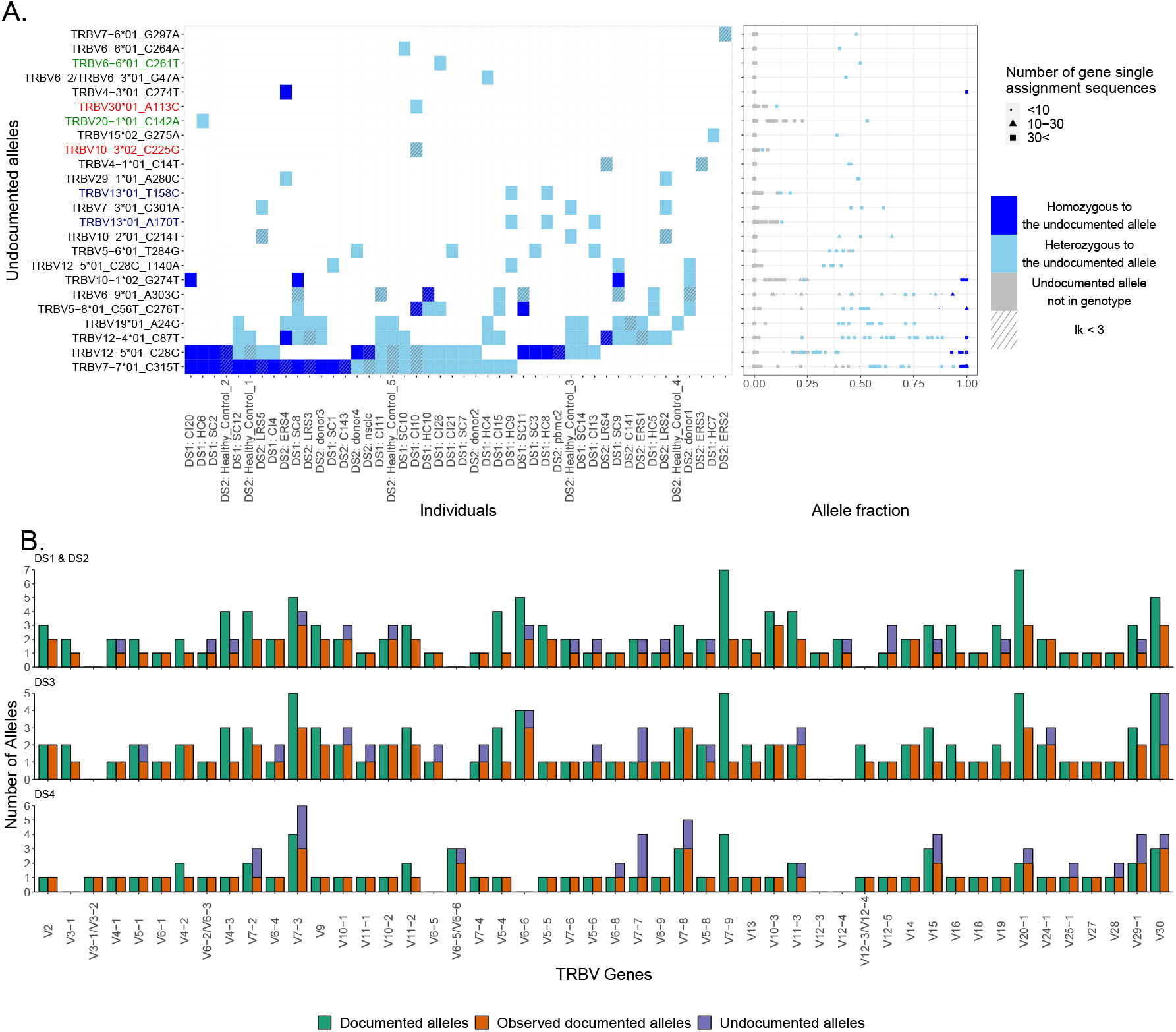
Undocumented alleles. (A) Undocumented allele distribution in DS1 and DS2. The left panel shows undocumented alleles within the genotype. Each row represents an undocumented allele, each column represents an individual. The Y-axis annotations in red, green, and blue correspond to alleles not following the expected multi-modal distribution, alleles adjacent nucleotide stretches, or both, respectively. The tile colors correspond to an individual’s genotype. The right panel shows the fraction of the undocumented allele assignments out of the gene assignments. The x-axis is the fraction and the Y-axis is the same as in (A). Colors correspond to an individual’s genotype of the allele. The shapes correspond to the number of gene assignments: a circle indicates that the number of gene assignments is less than 10, a triangle indicates that the number of gene assignments is between 10 and 30, and a square indicates that the number of gene assignments is more than 30. (B) Documented alleles versus the observed documented alleles and the undocumented alleles for each TRBV gene and in each data-set, DS1 and DS2, DS3, and DS4, respectively. The x-axis corresponds to the TRBV genes and the y-axis to the number of alleles. The color corresponds to the allele groups, dark red is the documented alleles, light red is the observed documented alleles, and blue is the observed undocumented alleles.

For further validation, we compared the undocumented alleles to three sources (Additional file 1:Table S5). In the first, an analysis of whole-genome short read sequencing data in 286 individuals from Luo et al. [11], six out of the 12 undocumented alleles were observed. The second was a pmTRIG [25] data-set, where we found four out of the 12. The third was a long read whole-genome sequencing data of 35 diploid donors from Ebert et al. [39] which confirmed six out of the 12. Altogether, eight of the 12 undocumented alleles were observed in at least one of these other data-sets, and two were observed in all of the data-sets. In addition to the 12 undocumented alleles, another five DS1 alleles matched known allele references in IMGT, but lacked 3’ nucleotides in IMGT (Additional file 1:Table S6). We replaced these IMGT alleles with their longer versions in our analyses, since it improves the genotype inference.

We also explored 5’ UTR genomic variation in DS1. Consensuses of the 5’ UTR were constructed as previously described in Mikocziova et al. [42, 43]. We compared the consensus sequences to the upstream regions, that include leader 1 (L-PART1), leader 2 (L-PART2), and the upstream leader 1 sequence (Additional file 1:Table S7), and found 31 undocumented upstream sequences. The L-PART1 and L-PART2 of 10 of these sequences are absent from IMGT, and the L-PART1 of TRBV10-3*02 is also absent from IMGT (Additional file 1:Table S8). Four of the 11 absent L-PART1 sequences and six of the 10 absent L-PART2 sequences were also observed in long-read assemblies (Additional file 1:Table S8). Two sequence variants were observed in L-PART2. Both were associated with TRBV13*01 and were also observed in the long-read assemblies. In L-PART1, we found 15 sequence variants from 14 different alleles, and 13 of them were also observed in the long-read assemblies. In addition, we found four alternative splicing sequences from three different alleles. The first two are from clusters of TRBV23-1, an ORF gene that lacks a functional splice donor site [44]. As a result, the 5’UTR consensus sequence of TRBV23-1*01 contains an intron between L-PART1 and L-PART2. Both consensus sequences differ from the previously reported upstream sequence by the number of copies of the TTTTG motif (Additional file 1:Fig. S3). The other two alternative splicing variants were found upstream of the TRBV7-7 alleles. Here, both the documented *01 allele and the undocumented allele *01_C315T carry the alternative splicing sequences. Three more upstream inferred sequences that correspond to TRBV4-3*01, TRBV20-1*01, and TRBV20-1*02 are absent from IMGT. However, those three sequences match the reference of the TRB locus under GRCh38 (Genome Reference Consortium Human Build 38) [40]. The 3’ ends of the L-PART1 references of the three sequences in IMGT seem to originate from the intron, and the 5’ splice site of the introns of those three alleles were likely misidentified in IMGT (Additional file 1:Fig. S4). We also found three variant consensus sequences associated with TRBV6-2*01/TRBV6-3*01 (Additional file 1:Fig. S3), all three were observed in the long-read assemblies for the TRBV6-2*01 annotation. One of them, TRBV6-2*01/TRBV6-3*01_2 was also observed in the undocumented allele TRBV6-3*01_G47A.

Next, we analyzed DS2, which includes data from 25 individuals. Fifteen undocumented alleles were inferred (Fig. 2A). Nine of them were also observed in DS1. Four others were present in the short-read whole genome databases. One was in Luo et al. [11] and all four were in pmTRIG [25]. Two out of those four undocumented alleles were observed in the 35 diploid long-read assemblies from Ebert et al. [39]. One out of those two undocumented alleles was observed in all sources, and two of the 15 undocumented alleles were not observed in any of the sources, including DS1 (Additional file 1:Table S5).

Lastly, we investigated genomic variation in DS3 and DS4. Both data-sets contain partial sequences, making it impossible to distinguish between alleles that differ in the regions outside those covered by the library primers. At first, we confirmed our inference approach by using artificial sequences trimmed from DS1 to the same sequence lengths as DS3 and DS4 (BIOMED2 and Adaptive, respectively; Additional file 1:Table S9). Of the 12 undocumented DS1 alleles, seven were in range for the BIOMED2 primers and two for the Adaptive primers. The artificial data sets enabled the inference of all the in-range undocumented alleles for both partial libraries. Having gained confidence in the approach, we then applied it to DS3 and DS4. From the 219 DS3 TRB genotypes [30], we inferred 29 potential undocumented allele patterns. However, 13 alleles failed the quality filtering steps (see Methods and Additional file 1:Table S10). Five of the remaining 16 undocumented alleles were independently identified in multiple genotypes, giving us added confidence in these inferences. Four alleles were supported by their identification by Luo et al. [11]. Six alleles were also supported by pmTRIG [25]. Eight alleles were not observed in any of the sources including DS1 and DS2 (Additional file 1:Table S10). A similar analysis of inferred genotypes from 786 individuals in the Adaptive Biotechnologies data-set (DS4) identified a further 24 undocumented alleles (Additional file 1:Table S11). We discarded 10 alleles that failed the quality filtering steps (see Methods and Additional file 1:Table S11). Six out of the remaining 14 undocumented alleles were independently identified in multiple genotypes, giving us added confidence in these inferences. Two alleles were supported by the presence of alleles identified also by Luo et al. [11]. Five alleles were supported also by pmTRIG [25]. Eight alleles were not observed in any of the sources including DS1, DS2, and DS3 (Additional file 1:Table S11). To summarize this part, we compared the documented alleles for this locus with all alleles observed in our data-sets and with the assemblies of genomic long reads (Fig. 2B). All in all we identified 39 undocumented TRBV alleles and 31 undocumented upstream sequences. Eight of those complete undocumented alleles and seven undocumented upstream sequences were also observed in the 35 long-read diploid assemblies of Ebert et al. [39].

### Inference of double chromosome deletions in four TRBV genes

Deletion polymorphisms can be so common that an individual may carry the deletion in both chromosomes. We refer to these genes as double chromosome deletions, although it is important to note that they are deleted from the expressed repertoire and not necessarily from the genome itself (see discussion). TRBV gene usage in DS1 shows such deletion polymorphisms in four genes (Fig. 3A). TRBV4-3 and TRBV3-2 are absent from the genotypes of eight individuals, while TRBV11-1 is absent from two individuals, and TRBV30 from one individual (Fig. 3B). In DS2, a similar data-set to DS1 from the point of view of sequence length of the coding region, double chromosome deletions of TRBV4-3 were identified in eight individuals, TRBV3-2 in two individuals, and TRBV7-3 in one individual (Additional file 1:Fig. S5). Interestingly, the individual from DS1 with an inferred TRBV30 deletion (HC10) was shown to be homozygous or hemizygous for the undocumented allele TRBV30*03_T285C. On the assumption that this undocumented allele is found at relatively low frequency within the human population, homozygosity is unlikely. However, it is possible that the undocumented allele has escaped more widespread detection because of its low usage level. This low usage is a consequence of TRBV30*03_T285C being a pseudogene, because its coding region includes an in-frame stop codon. HC10 therefore has at least the functional equivalent of a double chromosome deletion.

**Figure 3.**
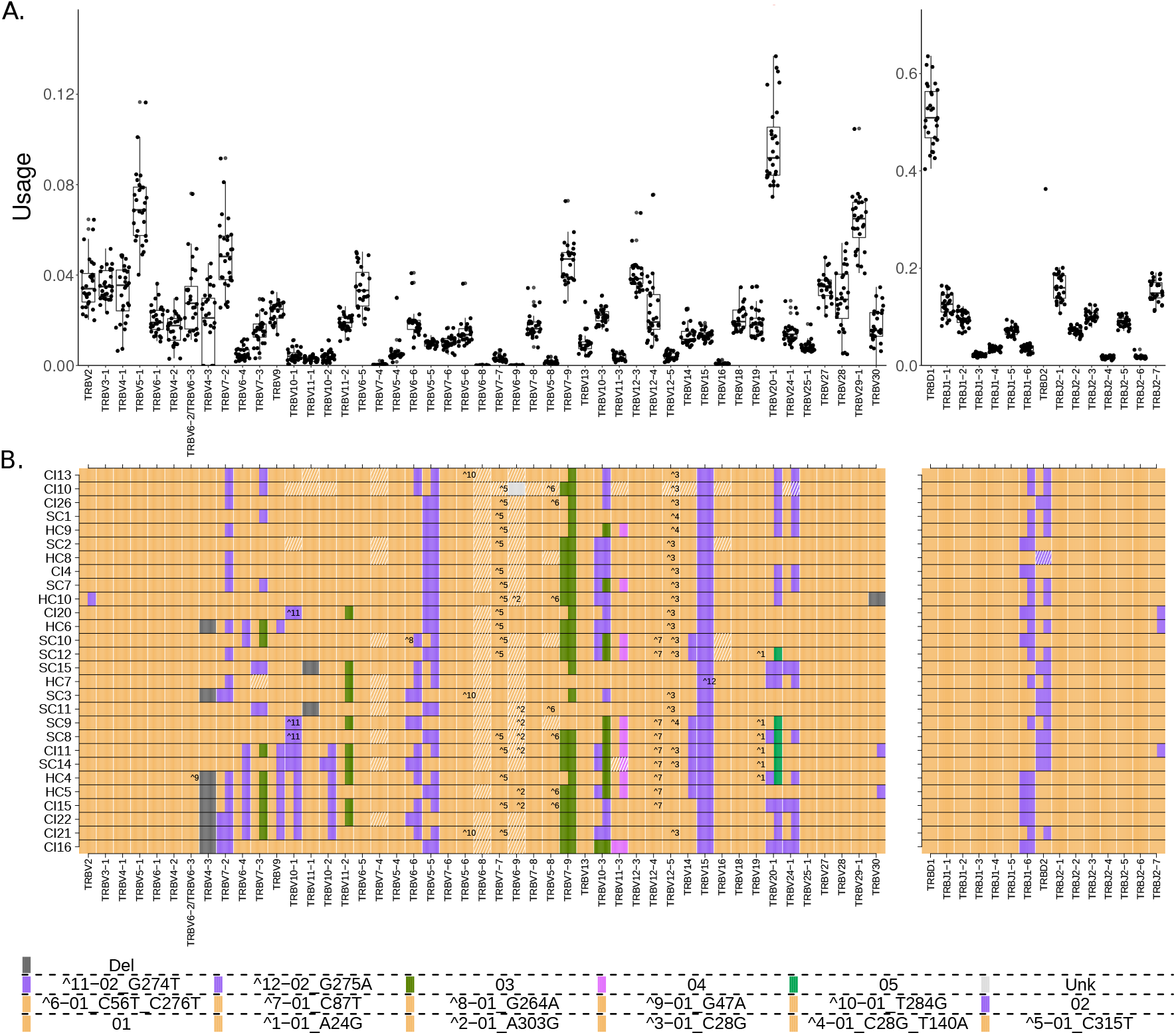
Gene usage and genotypes of the DS1 data-set. (A) TRB gene usage. Each dot is one individual’s relative gene usage, which is the fraction of the number of sequences mapped to a given gene by the number of total sequences. The X-axis shows the TRB genes in the order in which they are found in the genome, except TRBV30 that is located after the TRBD and TRBJ section. The Y-axis shows the frequency of gene usage. (B) TRB genotypes. Each row shows an individual genotype, and columns correspond to the different genes in the same order as in (A). Colors correspond to alleles as indicated in the bottom of the figure.

The gene TRBV4-3 and the pseudogene TRBV3-2 were always inferred as being deleted together in DS1 individuals (Fig. 3B). TRBV4-3 and TRBV3-2 are close to each other, and a common ~21Kb deletion which includes TRBV4-3, TRBV3-2 and one of the genes TRBV6-2/TRBV6-3 has been reported from genomic studies [10, 45-47]. The inference of a deletion of either TRBV6-2 or TRBV6-3 in AIRR-Seq data is made difficult, because the deletion might be hidden by the presence of the other identical TRBV6-32/TRBV6-2 gene. However, indirect evidence that the deletion polymorphisms seen in DS1 are associated with the previously-reported ~21Kb deletion comes from the usage of TRBV6-2*01/TRBV6-3*01. In individuals who lack TRBV4-3 and TRBV3-2, usage of TRBV6-2*01/TRBV6-3*01 is significantly lower than in the individuals who express TRBV4-3 and TRBV3-2 (Additional file 1:Fig. S6). It is therefore likely that detection of TRBV6-2*01/TRBV6-3*01 in these individuals is entirely a consequence of sequences utilizing one of the genes. This line of reasoning also allowed us to conclude that an undocumented polymorphism seen in sample HC4 is most likely an allele of the undeleted gene, meaning it can either be TRBV6-3*01_G47A or TRBV6-2*01_G47A.

In DS2 individuals, deletion of V4-3 was not always accompanied by evidence of deletion of TRBV3-2. This is likely because DS2 was collected from different sources, and in some data-sets non-productive sequences had been filtered out. Evidence of the presence or absence of the TRBV3-2 pseudogene is therefore lacking. In other samples, analysis of TRBV3-2 usage is compromised by its low usage (Fig. 3A).

### Strong asymmetry between the probabilities for misidentification of TRBD2 alleles

There are only two TRBD genes, TRBD1 and TRBD2, and three reported TRBD sequences (TRBD1, TRBD2*01 and TRBD2*02). Both genes are short and highly similar. TRBD1 is 12bp long, and TRBD2 is 16bp long. Each sequence includes a short central motif flanked by G-rich ends. A single G/A SNP that is flanked by runs of Gs differentiates the two TRBD2 alleles (TRBD2*01:”GGGACTAGCGGG**G**GGG”, TRBD2*02:”GGGACTAGCGGG**A**GGG”). During VDJ rearrangement, the ends of the TRBD segment are trimmed, and P-nucleotides and N-nucleotides are added between the joining TRBV, TRBD and TRBJ genes [48]. Studies of N-nucleotide addition in BCR V(D)J genes show the process to be biased towards addition of Gs, and addition of homopolymer tracts [49]. The unequivocal identification of germline-encoded nucleotides within the TCR*β* V(D)J junctions is therefore problematic, and this is particularly true for the TRBD gene ends. To reduce errors, we limited the TRBD gene genotype analyses to sequences with a minimum inferred length of nine bp. Some errors still remained, particularly as a result of TRBD2 allele assignment errors. Twenty three out of the 28 individuals from DS1 were initially inferred to be heterozygous at the TRBD locus. However, this is most unlikely according to the Hardy-Weinberg principle, which states that in equilibrium, in the absence of selection or other evolutionary pressures, the maximum frequency of heterozygous individuals in a population having two allelic variants of the gene is 0.5 [50].

To correct for these errors, we applied a process to determine the borders between TRBD2 genotype groups, as described in the methods (*Determining the borders between TRBD2 genotype groups*), resulting in a clear separation between the groups. This implies that TRBD2*01 is misidentified as TRBD2*02 with a probability of 4%. The probability of mis-identifying TRBD2*01 as TRBD2*02 is estimated to be 12.7%. We can thus estimate the average usage of TRBD2*01 in heterozygous individuals corrected for these errors to be ~ 0.393. There is a strong asymmetry between the probabilities for misidentification of TRBD2*01 as TRBD2*02 (GGGACTAGCGGG**G**GGG and GGGACTAGCGGG**A**GGG respectively) or the opposite. Misidentification is generally a result of several processes: 1. VDJ recombination, where the ends of the TRBD segment are trimmed, and P-nucleotides and N-nucleotides are added between the joining TRBV, TRBD, and TRBJ genes. ([49]) 2. PCR errors [51] 3. Sequencing errors [38, 52]. All three contribute to this asymmetric misidentification. It may also result from selection, from the amino acid differences in the sequences or from unknown structural variants in the locus that could be associated with the different alleles.

### Linkage disequilibrium between TRBJ1-6 and TRBD2

To explore J genotypes, we first checked for evidence of errors by exploring the fraction of all TRBJ1-6 assignments that are assigned to TRBJ1-6*01. This is the most likely error in TRBJ genotyping, as TRBJ1-6*01 is the only TRBJ gene with two known functional alleles. The distribution of frequencies shows a good partitioning between homozygous and heterozygous individuals (Additional file 1:Fig. S2D), indicating that the TRBJ1-6 alleles can be reliably inferred.

Of note, in heterozygous individuals TRBJ1-6*02 is considerably more frequently used compared with TRBJ1-6*01 (Additional file 1:Fig. S2D). The average fraction of TRBJ1-6*01 out of all sequences assigned to TRBJ1-6 in heterozygous individuals is ~ 0.39, which is comparable with the average fraction of TRBD2*01 out of all sequences assigned to TRBD2 in TRBD2 heterozygous individuals after correcting for mis-assignments (see above). The similarity between the biased usage of TRBJ1-6 and TRBD2 alleles in heterozygous individuals led us to test the genetic dependency between these loci.

The distance between TRBJ1-6 to TRBD2 is relatively short (~6000bp), suggesting these loci could indeed be in linkage disequilibrium (LD). To test this hypothesis, we reviewed Whole Genome Sequencing (WGS) records from the 1000 Genomes Project to profile the region’s variants based on TRBD2 haplotype (Additional file 1:Fig. S7). We observed SNPs with a high LD score between the genes. Further, the WGS haplotypes showed several other SNPs with high LD score scattered in the TRBD-TRBC2 genomic region, which strengthens the association between TRBD2 alleles and other markers in the locus. DS4 was unsuitable to test this hypothesis because DS4 sequences do not include the SNP that differentiates between TRBJ1-6*01 and TRBJ1-6*02. We therefore tested the LD hypothesis using DS3. Only genotypes for which we were confident of the TRBD2 genotype were taken into account. These genotypes are shown outside the gray areas of Additional file 1:Fig. S2B. Additional file 1:Fig. S8 shows that all of the homozygous TRBD2*01 individuals are also homozygous for TRBJ1-6*02. Also, 29 out of the 31 homozygous TRBJ1-6*01 individuals are homozygous for TRBD2*02.

### TRBJ usage is strongly dependent on the TRBD2 genotype

Having confirmed that TRBJ1-6 and TRBD2 are in LD, we next investigated the influence of TRBD2 genotypes on TRBJ/TRBV gene usage in the repertoires. Since such an investigation requires accurate TRBD2 genotype inference, accurate annotations of TRBJ genes, and a large data-set, DS4 was used.

We found that homozygous TRBD2*02 individuals tend to use TRBD2 1.29 times more than homozygous TRBD2*01 individuals (Fig. 4A, left panel). TRBD2 can undergo rearrangements only with TRBJ2 genes [53], so we expected that the usage of TRBJ2 genes should increase in homozygous TRBD2*02 individuals. Indeed, homozygous TRBD2*02 individuals use TRBJ2 genes significantly more than heterozygous and homozygous TRBD2*01 individuals (Fig. 4A, right panel), with 11 out of the 13 genes yielding p values lower than 0.001 (Mann-Whitney test, adjusted by Bonferroni correction). Furthermore, comparing the combined usage of all TRBJ2 genes to the combined usage of all TRBJ1 genes reveals a strong effect. For TRBD2*01 individuals, the mean usage of TRBJ1 was 0.473 compared to 0.366 for the TRBD2*02 individuals (Fig. 4B).

**Figure 4.**
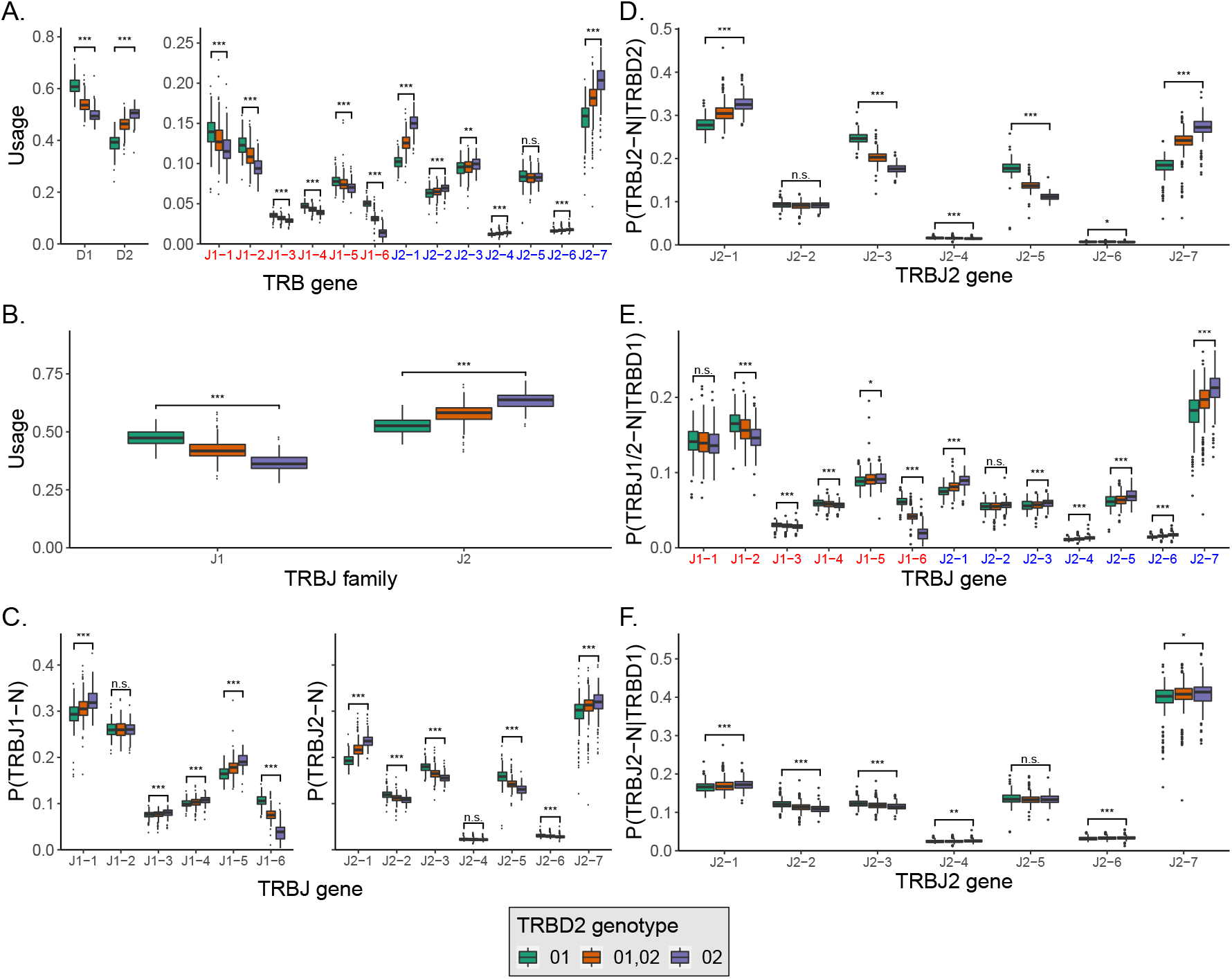
TRBD and TRBJ usage correspond to TRBD2 genotype. (A) TRBD and TRBJ gene usage in DS4 individuals with different TRBD2 genotypes. TRBJ genes are shown along the X-axis in the order in which they appear in the genome. (B) The TRBJ family usage in DS4 individuals with different TRBD2 genotypes. TRBJ families are shown along the X-axis in the order in which they appear in the genome. (C) TRBJ gene usage normalized to the TRBJ family usage, in DS4 individuals with different TRBD2 genotypes. TRBJ genes are shown along the X-axis in the order in which they appear in the genome. (D) The relative fraction of TRBJ2 genes out of the sequences that were assigned to TRBD2 and were longer than 7nt. (E) The fraction of TRBJ2 genes out of the sequences that were assigned to TRBD1 and were longer than 7nt. TRBJ genes are shown along the X-axis in the order in which they appear in the genome. (F) The fraction of TRBJ genes out of the sequences that were assigned to TRBD1 and were longer than 7nt. TRBJ genes are shown along the X-axis in the order in which they appear in the genome. The box colors correspond to the TRBD2 genotype. Statistical significance was determined using a Mann-Whitney test and adjusted by Bonferroni correction ( n.s. - not significant, * - p < 0.05, ** - p < 0.01, and *** - p < 0.001).

Next we examined if and how TRBD2 haplotypes affect the relative usage of individual TRBJ1 and TRBJ2 genes. For this, we plotted the relative usage of individual TRBJ1 and TRBJ2 genes normalized independently for each gene family (Fig. 4C). Surprisingly, we found that TRBJ usage within each family is also affected by the TRBD2 genotype. Since TRBD2 can rearrange only with TRBJ2 genes, we stratified the above distributions into subsets that include only biologically possible rearrangements. In particular, Fig. 4D shows the conditional probability P(TRBJ2-N|TRBD2) for all the TRBJ2 genes. The biased usage of the TRBJ2 genes observed in Fig. 4C is still present, indicating that TRBD2 relative likelihood to recombine with TRBJ2 genes is strongly affected by the TRBD2 genotype. We then explored the TRBJ gene fraction of the sequences assigned to TRBD1 (P(TRBJ1/2-N|TRBD1), Fig. 4E), and observed that TRBD2 genotype is associated with the TRBD1 likelihoods to rearrange with individual TRBJ genes. We further investigated the effect of TRBD2 genotype on the likelihoods to rearrange with individual TRBJ2 genes only, and the effect was mostly eliminated in terms of magnitude P(TRBJ2-N|TRBD1), Fig. 4F).

The strong biases observed in Fig.4A-E can result from amino acid alterations in the sequence, from non-coding regulatory variants, or from unknown structural variants in the locus that are associated with the different alleles. To discriminate between these options, we repeated the analysis for non-functional sequences that resulted from frame-shifts between the TRBV and TRBJ genes. Such non-functional sequences are commonly used to reflect the initial V(D)J usage prior to thymus selection [54–56]. Of note, these non-functional clones cannot overlap with functional clones, since in T cells clones are defined as sharing identical VDJ sequence. In these non-functional sequences the biases are pronounced in a similar fashion (Additional file 1:Fig. S9). Thus, we conclude that the differences between the TRBD and TRBJ rearrangements stratified by TRBD2 genotype are most likely due to structural differences or non-coding regulatory variants between the haplotypes rather than due to negative selection.

### Haplotype inference reveals several association patterns

From the genotype analysis of DS1, we observed a potential pattern between homozygosity of TRBV7-2*02 and lack of usage of TRBV4-3 in four individuals. To inspect the link between TRBV7-2*02 and the lack of usage of TRBV4-3, we turned to haplotype inference. In an analogous way to the haplotyping methods for B cell receptor data [21, 22], a heterozygous TRBD or TRBJ gene was needed as an anchor for TRBV haplotype inference. A suitable candidate is TRBJ1-6, for which 11 individuals from DS1 (Fig. 3B) and eight individuals from DS2 (Additional file 1:Fig. S5) are heterozygous. In seven individuals who are heterozygous for TRBV7-2, the chromosome that carried the allele 02 of this gene had a clear deletion of TRBV4-3 (SAMPLE_ANCHOR-GENE_ALLELE: CI13_J1-6_01, SC12_J1-6_02, HC6_J1-6_01, HC9_J1-6_02, SC7_J1-6_02, LRS2_J1-6_01, and donor3_J1-6_02). The genotype and haplotype inferences both support the association between TRBV7-2 genotype and the usage of TRBV4-3. To further quantify this effect, we surveyed individual genotype and haplotype inferences in DS3. DS3 is a much larger data-set, which covers one of the unique SNPs of TRBV7-2*02, allowing the differentiation of TRBV7-2*02 from the rest of the known TRBV7-2 alleles (Additional file 1:Table S2). In eight out of nine individuals with a high genotype inference likelihood, the pattern between homozygosity of TRBV7-2*bp02 (see section *Allele pattern collapsing*) and a deletion inference of TRBV4-3 was apparent (Additional file 1:Fig. S10 and S11). Another gene with a link to TRBV7-2 is TRBV6-2/TRBV6-3. Its usage was also highly affected by the genotype of TRBV7-2. The mean usage of TRBV6-2/TRBV6-3 in TRBV7-2*bp02 homozygous individuals was less than half of the mean usage in TRBV7-2*bp01 homozygous individuals. Since we cannot distinguish between TRBV6-2 and TRBV6-3, this observation supports the hypothesis that both TRBV4-3 and one of TRBV6-2/TRBV6-3, are not present in haplotypes that carry TRBV7-2*bp02. We used long-read assembly haplotypes to investigate the connection between TRBV7-2 and the deletions. All 40 haplotypes that carried TRBV7-2*bp02 had a deletion stretch at the genomic location of TRBV4-3 and either TRBV6-2 or TRBV6-3. An additional 27 haplotypes carried TRBV7-2*bp01, of which eight had the same deletion stretch as with TRBV7-2*bp02.

In addition, the following association patterns between specific alleles and single chromosome deletions were revealed by haplotype inference: 1. TRBV24-1*02 was observed in eight samples on the chromosome carrying TRBJ1-6*02 and none in the other chromosome. 2. TRBV28*Del was observed in 4 samples on the chromosome carrying TRBJ1-6*02 and none in the other chromosome. 3. TRBV20-1*05 was observed in five samples on the chromosome carrying TRBJ1-6*01 and none in the other chromosome. 4. In eight individuals when TRBV24-1*02 was present on chromosome TRBJ1-6*02, TRBV20-1*02 was also observed. Of note is a haplotype block between the genes TRBV6-4 to TRBV10-1, that was observed in DS1:CI21 on the TRBJ1-6*01 chromosome and in DS2:donor4 on the TRBJ1-6*02 chromosome (Fig. 5).

**Figure 5.**
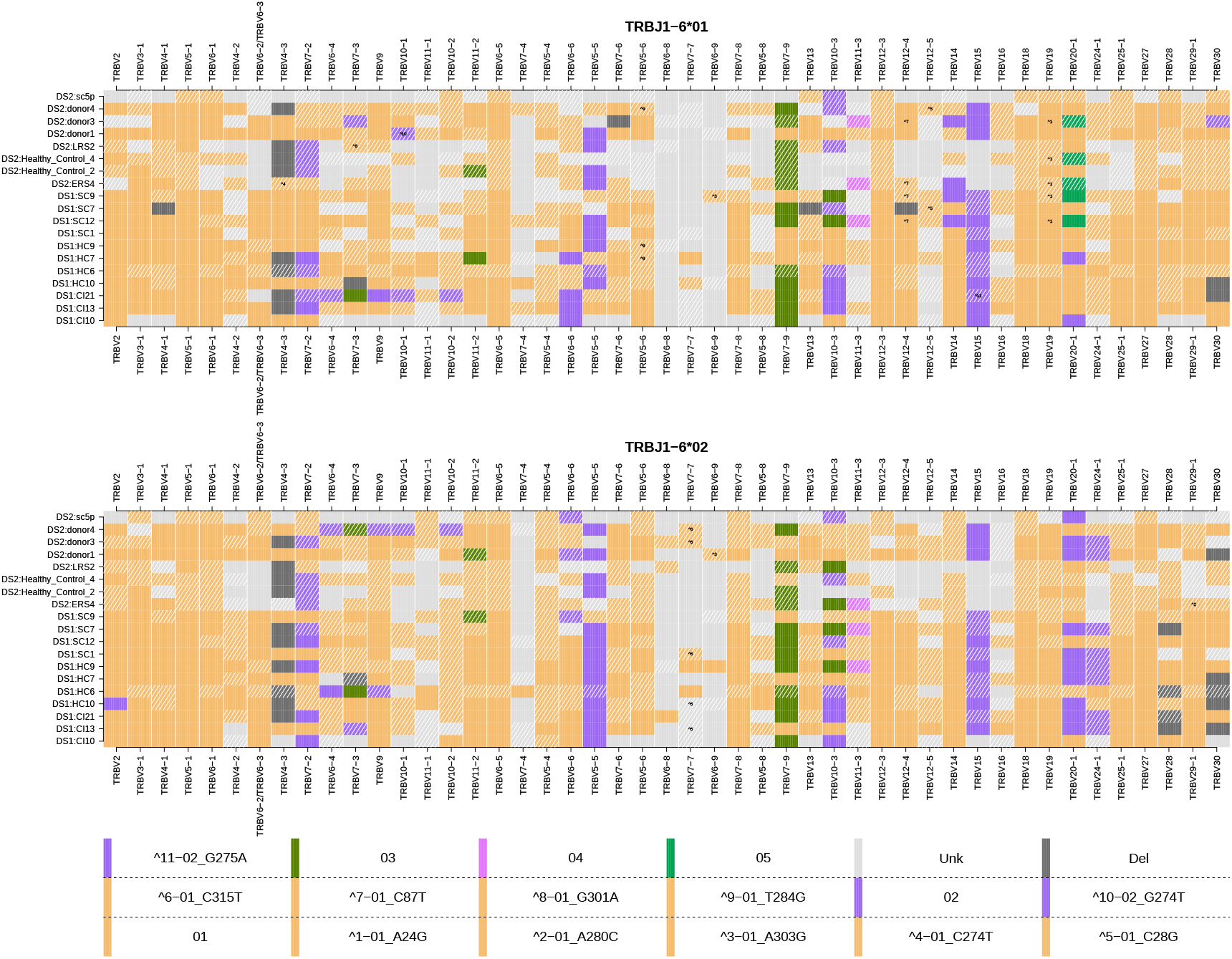
TRBV haplotypes for 19 individuals from DS1 and DS2. The upper and lower panels show the TRBV haplotypes anchored with TRBJ1-6*01 and TRBJ1-6*02, respectively. Each row is an individual’s haplotype, and each column is a V gene call. The colors correspond to the V alleles and the tile annotations correspond to the undocumented allelic variants.

TRBD2 was also considered a potential anchor gene for haplotype inference. To examine the reliability of this inference, we used the same individuals who were heterozygous for TRBJ1-6. Ten out of the 11 individuals from DS1 were heterozygous for TRBD2 and were used for the haplotype inference (Additional file 1:Fig. S12). Although the number of recombinations of TRBD2 with the TRBV genes is much larger than TRBJ1-6 and could potentially supply a better inference, comparison of the results from both anchor genes shows a different picture. The haplotypes inferred with TRBD2 commonly show occurrences of more than one allele per gene on a single chromosome. This is most likely due to ambiguous assignment of the very short and similar TRBD2 alleles. Hence, although haplotype inference with TRBD2 is feasible, it is likely to be less accurate.

## Discussion

In this study, we have explored how beta chain TCR VDJ sequences of different lengths, generated using differing technologies, can contribute to our knowledge of TRB genes, their allelic variants, and their influence on the expressed repertoire. Previous studies of BCR genotypes and haplotypes have led to the identification of dozens of new allelic variants of heavy and light chain variable region genes. This approach has not been extended to investigations of the TCR genes. Zhang et al. [57] investigated the reliability of V gene identification for different lengths of sequences of IGH, IGK, IGL, TRA, and TRB with respect to somatic hypermutations and sequencing errors. The comparison there is between full-length and lengths of 100, 150 or 200 nucleotides, whereas we used an analogous approach to compare the lengths of the common sequencing protocols. The focus of most TCR gene studies remains firmly fixed on the CDR3 regions of the genes [58]. Despite their interaction with the Major Histocompatibility Complex (MHC), and their documented influences on TCR/MHC/peptide interactions [59, 60], the CDR1 and CDR2 gene sequences and their translated products are still generally ignored in TCR repertoire studies. Only the 3’ ends of the TCR variable gene sequences are included in the amplicons generated by commercial providers of TCR sequencing such as BGI, iRepertoire and Adaptive Biotechnologies. Their sequencing setup allows the unambiguous identification of most variable region genes that may partially encode the CDR3 sequences, but they are rarely able to identify TRBV genes at the allele level. This study sheds light on understudied regions of the TCR, to enable accurate identification of new alleles, genotypes and haplotypes.

Despite the intimate partnership of MHC proteins and TCRs in the recognition of antigenic peptides, it is only the genes of the MHC that are widely recognised as disease susceptibility genes [61]. The germline genes that rearrange to produce TCRs are rarely accorded much importance. The multiple sets of TCR alpha, beta, gamma and delta V, D and J genes each include many highly similar genes. This may have encouraged the view that the astonishing processes of V(D)J recombination should generate much the same kind of repertoire, no matter which germline genes are available to an individual.

Germline TCR genes may also represent a blind-spot to the immunological community, because until recently they were so difficult to document in an individual, let alone in a population. High throughput sequencing studies of TCR repertoires now enable easy documentation of germline genes. In this study we adapt tools and techniques that were developed for analysis of BCR genotypes [18, 19, 21, 23, 62, 63] and haplotypes [17, 22] for the analysis of TCR data-sets. Other tools such as LymAnalyzer have been specifically developed for the analysis of TCR data [64], but they do not produce genotypes and haplotypes.

Our analysis demonstrates that many undocumented V genes remain to be discovered, and that even short read sequences can be analysed for the detection of undocumented polymorphisms. It is clear, however, that much additional information is to be gained by the study of full length VDJ genes. Analysis of genotypes and haplotypes from full-length sequence data-sets of 53 individuals led to the identification of 18 TRBV alleles that are not documented in the IMGT Reference Directory. In contrast, only 16 polymorphisms were identified in truncated sequences generated from 219 individuals using BIOMED-2 primers, and just 14 new allele patterns were seen in the 786 Adaptive Biotechnologies data-sets. This latter result reflects both the very short lengths of the Adaptive Biotechnologies sequences, and the general lack of variability in the 3’ ends of the TRBV genes. To try and speculate whether the TRB polymorphisms are more important for MHC binding or antigen binding, we compared the distributions of known SNPs in TRBV and IGHV genes (Additional file 1:Fig. S13 and Fig. S14), expecting to find more amino acid replacements in the regions of CDR1/2. Schwartz et al. [65] have shown that germline diversity at a given position is a good indicator for the potential to survive after a somatic mutation at that position. However, there are far fewer known SNPs in TRBV than in IGHV, which makes it impossible to address this interesting question. We also could not address the question of polymorphisms in TRA, as there are very few full length TRA samples available.

It is interesting to contrast this level of gene discovery with genetic variation amongst the BCR-encoding genes: to date, 120 functional alleles are documented in IMGT for 48 TRBV genes (avg. 2.5 alleles per gene). This compares to 286 functional alleles reported for the 55 IGHV genes (avg. 5.2 alleles per gene). This study adds a total of 39 alleles and SNPs to the record of TRBV genes (~ 33% of the known alleles), and it seems likely that many undocumented TRBV alleles remain to be discovered.

It appears that there is less structural variation in the TRBV locus than is seen in the IGHV locus. A well-documented 21Kb deletion polymorphism in the TRBV locus, involving the TRBV4-3, TRBV3-2 and TRBV6-2 genes, was frequently noted here. Other deletion polymorphisms, each involving single genes - TRBV11-1, TRBV7-3, TRBV30, TRBV2, TRBV4-2, TRBV6-5, TRBV5-5, TRBV13, and TRBV28 - were seen at relatively low frequencies. No deletion polymorphisms involving the TRBD or TRBJ loci were detected. In contrast, numerous relatively common deletions are now known in the IGH loci, including one involving 13 consecutive, functional IGHV genes, and one involving six consecutive IGHD genes [21, 23]. The seven functional IGHV genes that can be found between IGHV4-28 and IGHV4-34 are rarely all present, or all absent, with at least six recognized structural variant haplotypes [16, 23].

It should be emphasised that the deletions reported here reflect an absence of rearrangements in the expressed TCR repertoire. It is possible that the deletions are ‘functional’ rather than structural, perhaps as a result of variants in recombination signal sequences (RSS) or other regulatory elements. Certainly in a few individuals, a handful of rearrangements of the genes in question were seen. For example, TRBV4-3 was usually present in about 2% of all rearrangements, but in eight individuals it was seen at frequencies less than 0.05

No previously unknown gene duplications were inferred in this study. This is in sharp contrast to observations from studies of the IGHV locus. Early RFLP (restriction fragment length polymorphism)-based analyses pointed to duplications of sequences that came to be known as IGHV3-23 and IGHV1-69 [66-70]. More recently, the duplication of these two genes and of three other functional genes were confirmed, first by analysis of AIRR-Seq data [71] and then from genomic assembly data, including the second complete assembly of the IGHV locus [14].

It was not possible in this study to explore possible RSS variants, as RSS are lost from the genome during VDJ recombination. On the other hand, 5’ RACE data allows for the exploration of variation in the 5’ UTR, as has recently been reported from BCR repertoire studies [42, 43]. Thirty one variants of the 5’ UTR sequences of TRBV genes were identified in this study – a similar level of variability to that seen in IGHV studies. The functional implications of this kind of variation is yet to be determined.

The documentation of allelic variation and structural variation in the TCR gene loci will be important, as there are clear consequences of such variation on the expressed TCR repertoire, and if such variation can be shown to have consequences for the disease susceptibility of different individuals. It might be assumed that thymic selection would so powerfully shape individual TCR repertoires that any consequences upon the TCR repertoire of individual genotypes and haplotypes would be obscured. This study shows, however, that the usage of particular genes in the expressed repertoire appears to be very similar between individuals. The ‘shape’ of the TCR repertoire may therefore be as predictable as has been found for the BCR repertoire [3, 4], reflecting both the carriage of individual genes and the LD that is found within the loci. Conspicuous LD identified in this study includes that of the TRBV4-3/TRBV3-2/(TRBV6-2 or TRBV6-3) deletion polymorphism and carriage of the TRBV7-2*02 allele, as well as linkage between the TRBD2 and TRBJ1-6 loci. These different haplotypes, in turn, are associated with significant differences in the usage of neighbouring genes.

The power of AIRR-Seq analysis is well demonstrated by the TRBD gene analysis in this study. Although most TCR AIRR-Seq studies have been CDR3-focused, the TRBD genes that provide recurring central motifs to the CDR3 have usually been ignored. This study demonstrates that meaningful analysis of TRBD genes is possible, and that even the slight sequence variation between the TRBD genes have consequences for the expressed repertoire. Within large VDJ data-sets, TRBD genes can be identified with confidence, and even the presence of different TRBD2 alleles in an individual’s genotype can shape the expressed repertoire in predictable ways. Yet, all of these findings were procured mainly from individuals from developed countries or undocumented geographic origin. Sequencing individuals from developing and least developed countries could very well add or alter some of these results, as argued by Peng et al. [72]. Nonetheless, as our knowledge of these and other genes of the TCR loci grows, the way will be open to identify ‘departures from the repertoire norm’ that may have biological and perhaps even clinical implications.

The frequency distribution analysis shown in Fig. 2A focuses on bi-or tri-modal distributions. In cases that involve duplicated genes that share the same allele, such as TRBV6-2 and TRBV6-3, having more than two alleles per gene may shift the location of the second/third/fourth mode of the distribution, but it will still not be unimodal. As such, as long as we do not have extreme cases in which the relative frequency of the candidate allele is comparable to the mis-identification rate of the allele, the method is applicable. In other cases, such as the ones present in the IGH/IGK loci, these assumptions should be revisited [73].

To date, there have been very few associations of TCR genes with disease. One clear example is the genetic predisposition to carbamazepine-induced Stevens-Johnson syndrome (SJS), a severe cutaneous hypersensitivity with high mortality [74]. SJS and other cutaneous hypersensitivity reactions have been linked to HLA types, but these associations have all had a low positive predictive value [75], leading others to explore a possible role for TCR genes. It has now been shown that SJS is associated with the usage by cytotoxic T cells of a public TCR clonotype encoded by the TRBV12-4 and TRBJ2-2 genes [76]. Full length sequences were not reported in the study of Pan and colleagues [76], and so a possible role for specific allelic variants of these genes could not be explored. Interestingly, in the present study a previously undocumented polymorphism of TRBV12-4 was identified.

The lack of disease associations with TCR genes is likely to be a reflection of our ignorance of individual genetic variation within the TCR loci. Only after thorough exploration of the population genetics of the TCR genes, and of individual variation in the expressed TCR repertoire, will it be possible to determine whether or not these genes have a role in disease susceptibility. Genomic sequencing of TCR genes will contribute to this [77], but the present study demonstrates that it will also be possible to do this efficiently through the analysis of AIRR-Seq data. For this reason, the amplification of full-length TCR V(D)J sequences, or genomic long read paired sequencing of the locus, must be strongly encouraged in AIRR-Seq studies. The results presented in this paper pave the way towards establishing functional links between the TRB germline repertoire and the TCR immune response. They expand our knowledge of genomic variation in the TRB locus and lay the ground for more accurate basic and clinical studies.

## Conclusions

To summarize our findings, we identified 39 undocumented TRBV alleles and 31 undocumented upstream sequences, and inferred double chromosome deletions in TRBV4-3, TRBV3-2, TRBV11-1, and TRBV30. For TRBD and TRBJ, we determined the error rates in identification between TRBD2*01 and TRBD2*02 and corrected them, discovered LD between TRBJ1-6 and TRBD2, and found a strong bias in the usage of TRBJ genes depending on which TRBD2 allele is used. For example, for TRBD2*01 individuals, the mean usage of TRBJ1 was 0.473 compared to 0.366 for the TRBD2*02 individuals. Overall, our study sheds light on the genomic loci encoding TRB, to enable identification of new alleles, genotypes and haplotypes.

## Declarations

### Ethics approval and consent to participate

Not applicable.

### Consent for publication

Not applicable.

### Availability of data and materials

Alleles inferred in this study are listed in Supplementary Data. Repertoire analyses of the four data-sets, the sequences of the undocumented alleles inferred, the IMGT TRBV references used, and the code used for the analyses are available in the Github repository (https://github.com/omaviv/TCR_genotype) [78]. The genotypes and haplotypes are also accessible through VDJbase [79] (https://www.vdjbase.org)

### Authors’ contributions

GY conceived the idea, analyzed the data, and supervised the work. AO and AP developed the methods and analyzed and interpreted the data. OLR and CTW analyzed the genomic data. PP, AO, AP, AMC, WL, and GY wrote the paper. All authors read and approved the final manuscript.

## Acknowledgements

We would like to thank Mats Ohlin, Martin Corcoran, and James Heather for stimulating discussions.

## Additional Files

Additional file 1 — Supplementary materials: TRB germline variability is revealed by inference from repertoire data

