## Supplementary material for "T Cell Receptor Beta Germline Variability is Revealed by Inference From Repertoire Data"

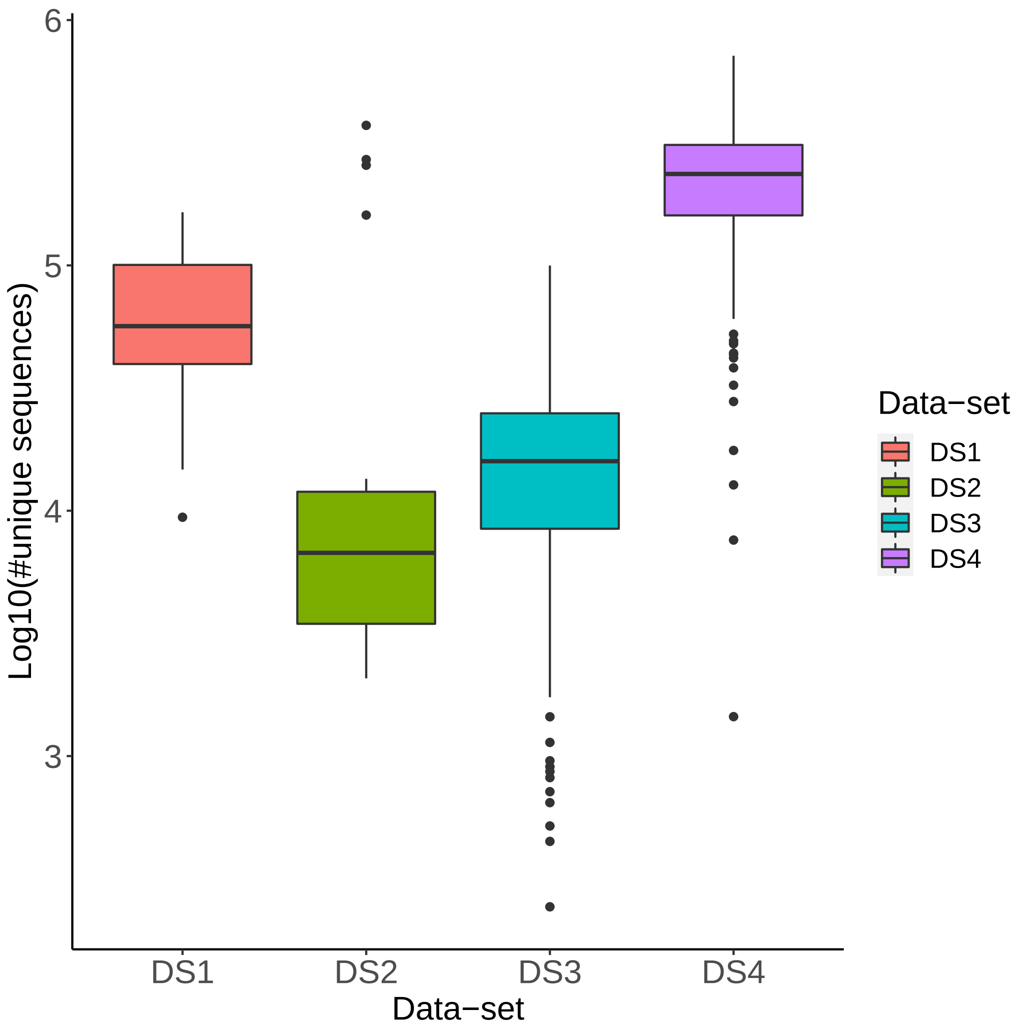


Fig S1: **Unique sequence distribution in the four data-sets** The X-axis represents the four data-sets. The Y-axis is the number of unique sequences by log10.

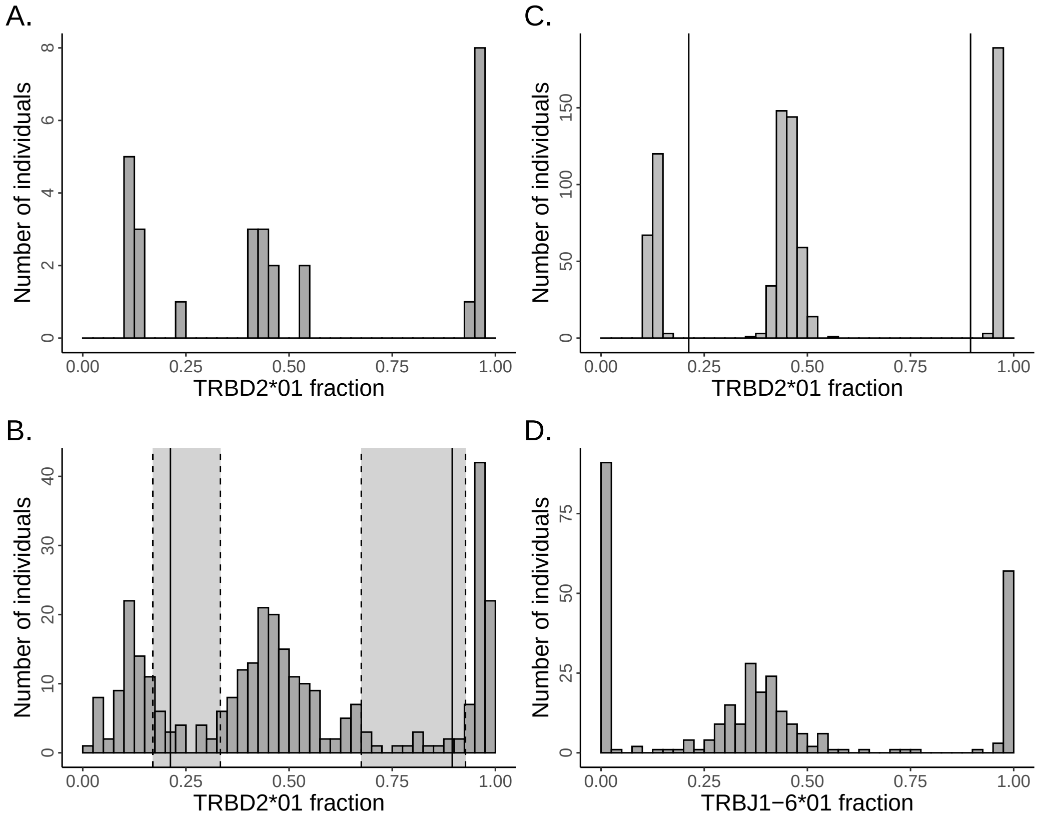


Fig S2: **TRBD2 and TRBJ1-6 genotypes and gene usage frequencies.** (A) The frequency of TRBD2*01 usage, as a fraction of total D2 usage in 28 individuals from DS1. Homozygous and heterozygous genotypes can be inferred from the distribution. (B) The frequency of TRBD2*01 usage, as a fraction of total D2 usage in 313 individuals from DS3. Solid vertical lines indicate boundaries between homozygous and heterozygous individuals, as calculated from analysis of DS4. The genotypes of individuals whose frequencies fall within the shaded regions cannot be inferred with confidence. (C) The frequency of TRBD2*01 usage, as a fraction of Total TRBD2 usage in 786 individuals from DS4. Solid vertical lines indicate boundaries between homozygous and heterozygous individuals. (D) The frequency of TRBJ1-6*01 usage, as a fraction of Total TRBJ1-6 usage in 313 individuals from DS3.

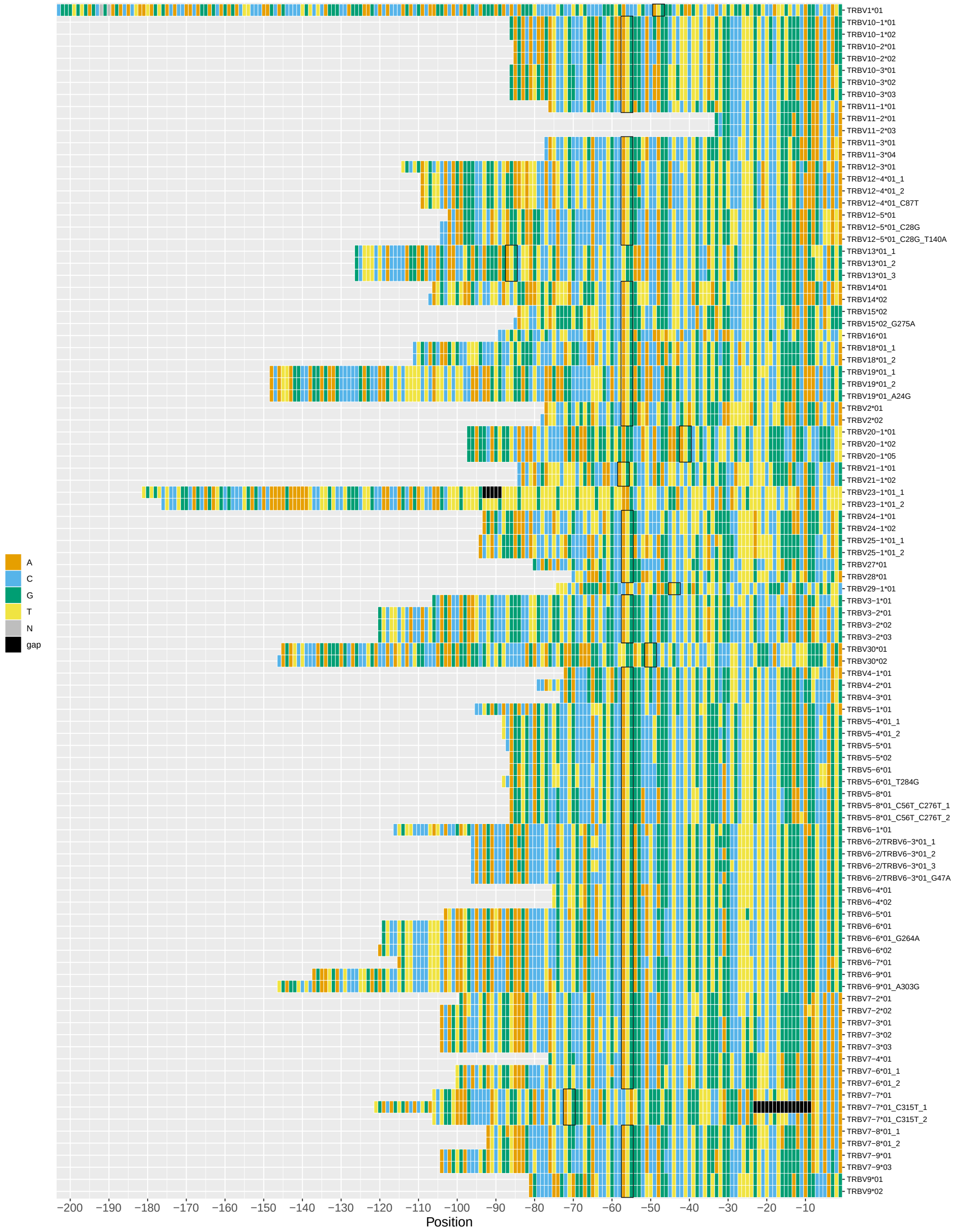


Fig S3: **5′ UTR nucleotide sequences of TRBV genes.** Each row is a consensus sequence for an allele. The column positions are numbered from the start of the FR1 region. Start codons are indicated with black frames.

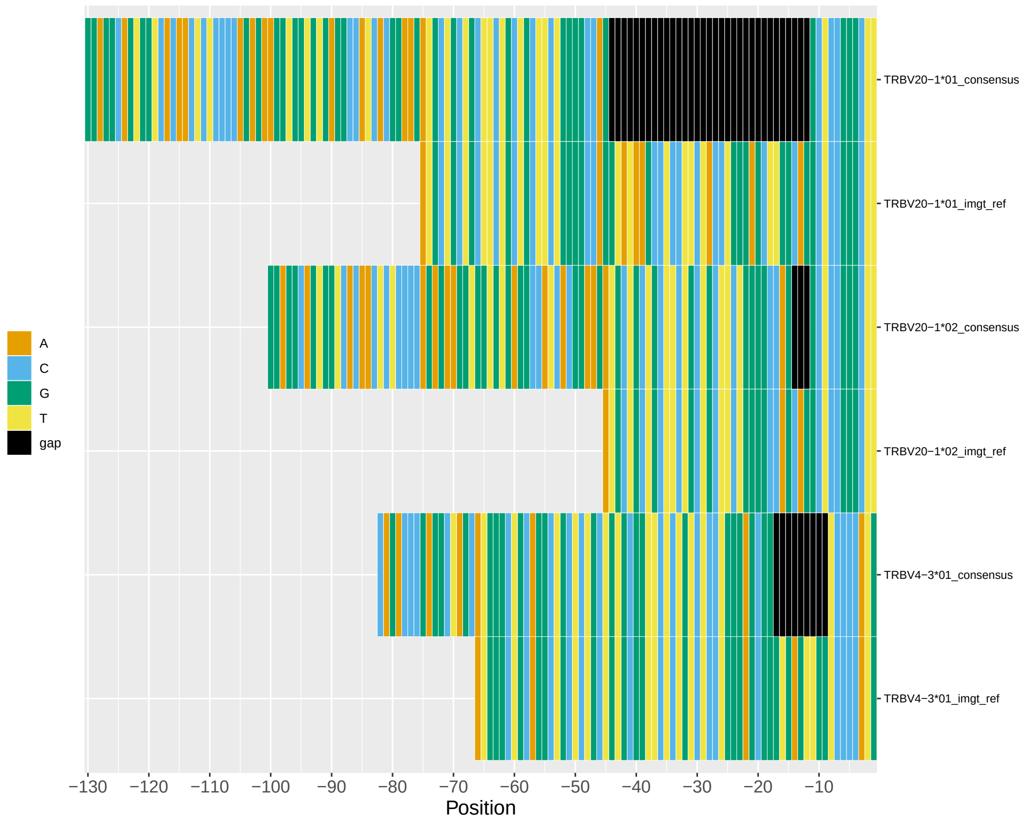


Fig S4: **5′ UTR nucleotide sequences of TRBV genes.** Each row is or a consensus sequence for an allele or the IMGT reference sequence for the allele. The column positions are numbered from the start of the FR1 region. The gaps for the consensus sequences were opened between the end of the L-PART1 to the beginning of the L-PART2.

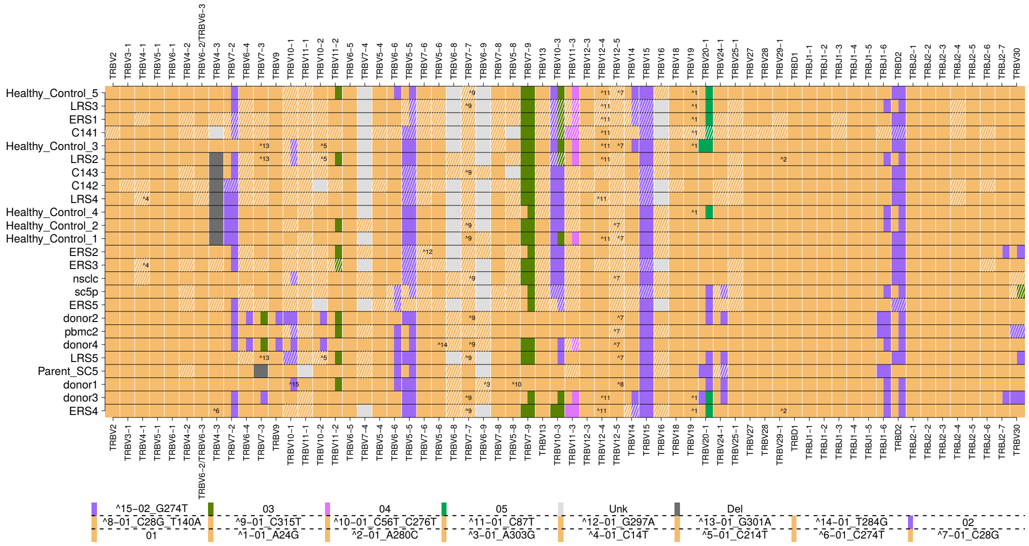


Fig S5: **DS2 TRB genotype heatmap.** Each row is an individual, each column is a gene sorted by locus. Colors correspond to alleles.

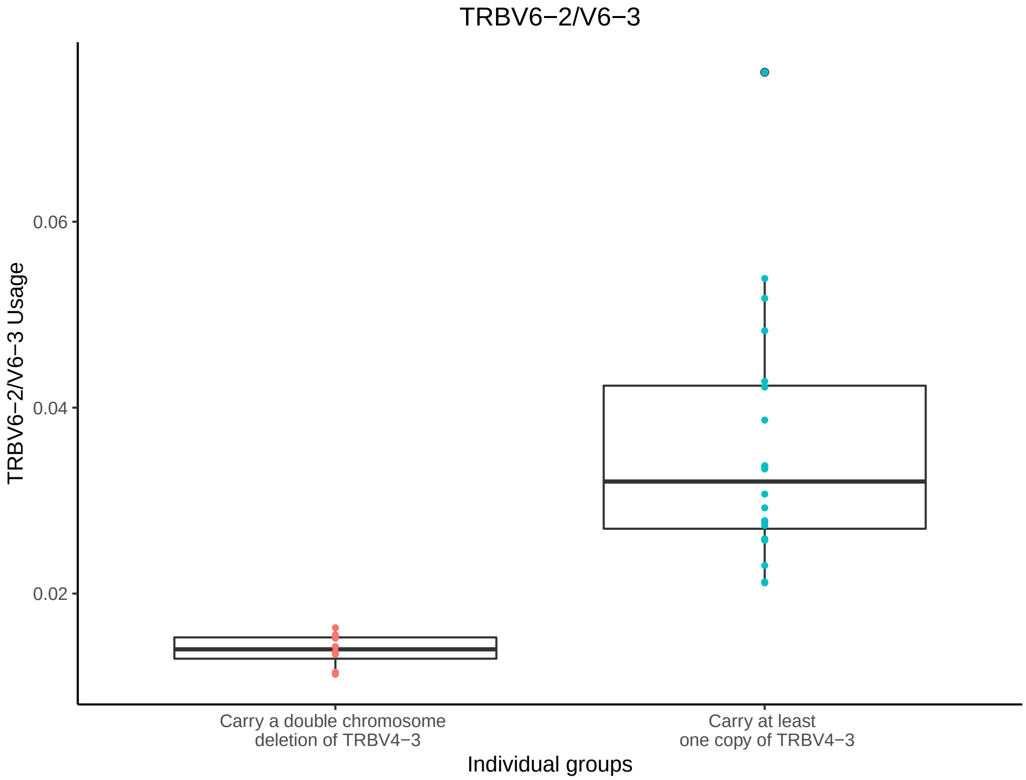


Fig S6: **TRBV6-2/TRBV6-3 usage correlates to the existence of TRBV4-3 and TRBV3-2 in DS1.** The X-axis represents the two groups of individuals, the Y-axis is the TRBV6-2/TRBV6-3 usage out of all the TRBV genes, and the colors correspond to the individual group.

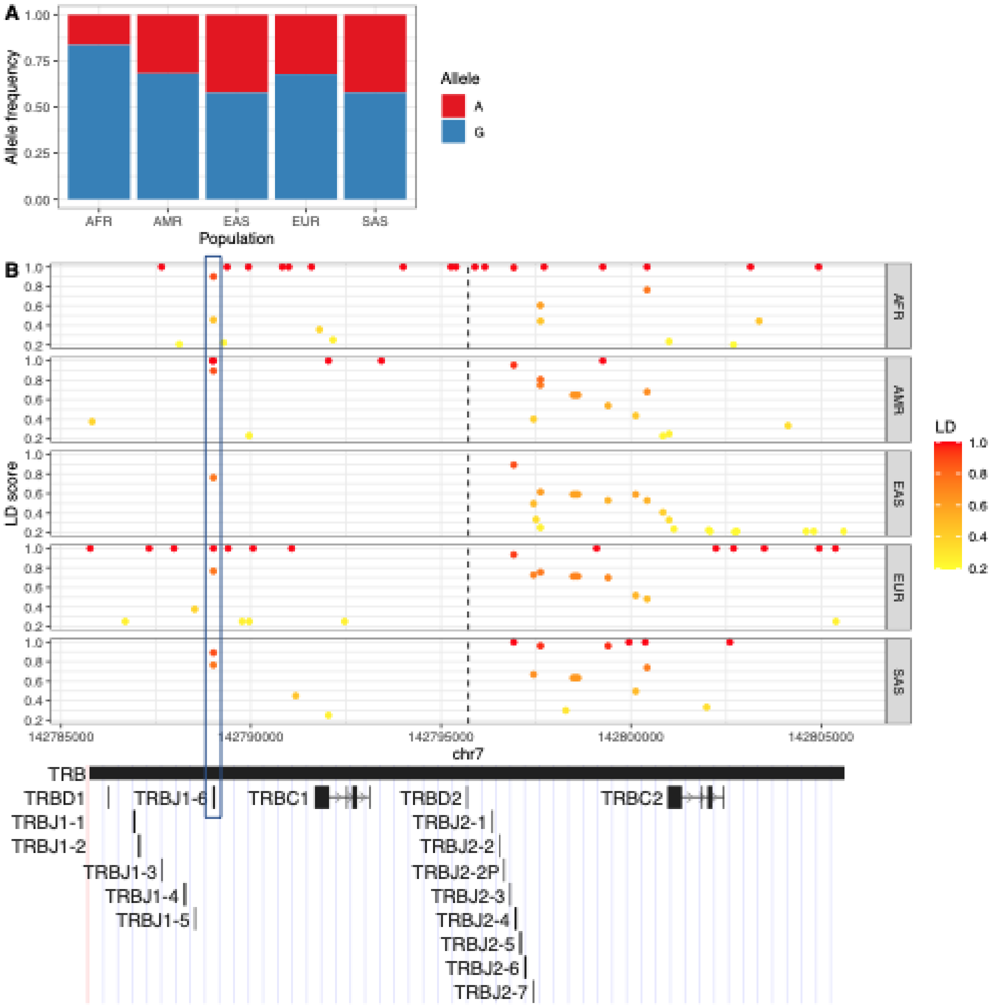


Fig S7: **Linkage disequilibrium in TRBD2 haplotype.** Whole genome sequencing was used to haplotype, based on TRBD2 alleles as anchors, and pinpoint potential linkage disequilibrium SNPs within the TRBD-TRBC2 chr7 region. (A) The allelic frequency of TRBD2 within each population. The Y-axis is the allele frequency, the x-axis is the different population groups. The colors represent the allele, blue for G and red for A. (B) Genome data viewer for the TRBD-TRBC2 chr7 region. The upper panel shows the LD score of each variant per position and for each population group. Each row is a population group, and each column is a genomic position in chr7. Y-axis is the LD score, and each dot is a genomic variant. The color scale represents the LD score value. The lower panel shows the genomic annotations for the selected chr7 region. The annotations' locations correspond to the upper panel x-axis genomic positions. The dashed line points to the SNP position that differentiates between the TRBD2 alleles, and the rectangle marks TRBJ1-6.

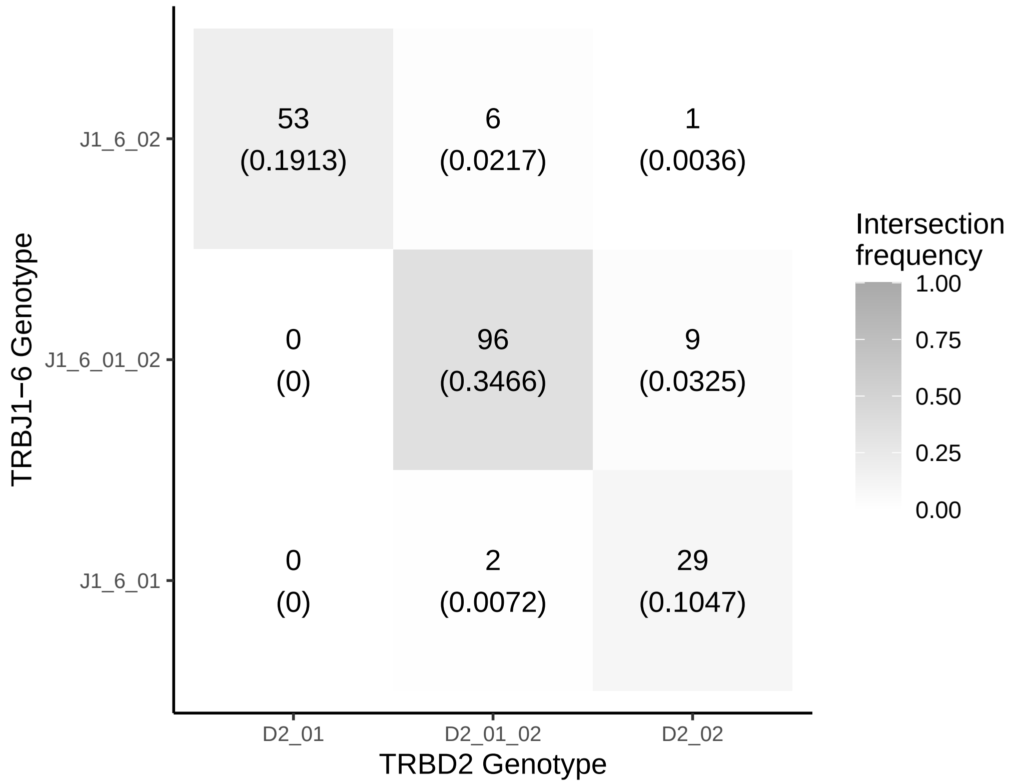


Fig S8: **Genomic correlation between TRBD2 and TRBJ1-6 in DS3.** The number and proportions of individuals in DS3 as observed with different TRBJ1-6 and TRBD2 genotypes.

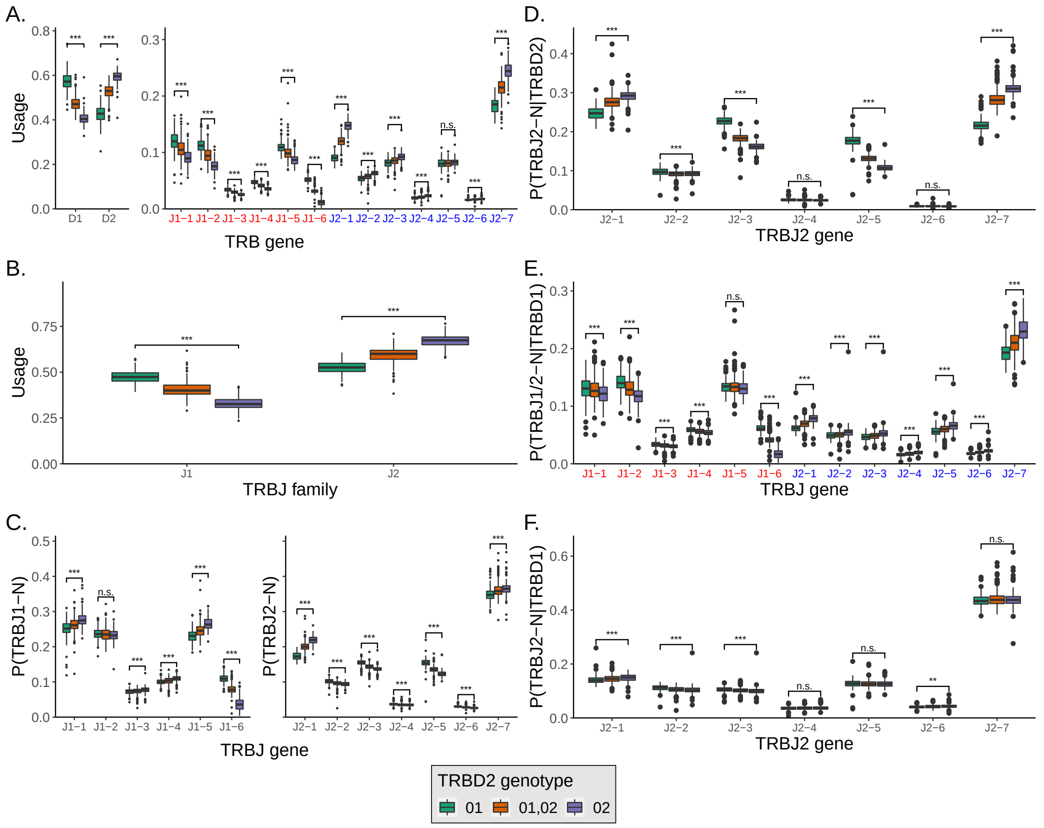


Fig S9: **TRBD and TRBJ usage out of the non-functional sequences corresponds to TRBD2 genotype in DS4.** (A) TRBD and TRBJ gene usage in DS4 individuals with different TRBD2 genotype. TRBJ genes are shown along the X-axis in the order in which they are found in the genome. (B) The TRBJ family usage in DS4 individuals with different TRBD2 genotypes. TRBJ families are shown along the X-axis in the order in which they appear in the genome. (C) TRBJ gene usage normalized according to the TRBJ "family" usage in DS4 individuals with different TRBD2 genotype. TRBJ genes are shown along the X-axis in the order in which the genes are found in the genome. (D) The fraction of TRBJ2 genes out of the sequences that were assigned to TRBD2 and were longer than 7nt. (E) The fraction of TRBJ2 genes out of the sequences that were assigned to TRBD1 and were longer than 7nt. TRBJ genes are shown along the X-axis in the order in which they are found in the genome. (F) The fraction of TRBJ genes out of the sequences that were assigned to TRBD1 and were longer than 7nt. TRBJ genes are shown along the X-axis in the order in which they are found in the genome. The boxes' colors correspond to the TRBD2 genotype. Statistical significance was determined using a Mann-Whitney test and adjusted by Bonferroni correction ( n.s. - not significant, * - $p < 0.05$, ** - $p < 0.01$, and *** - $p < 0.001$).

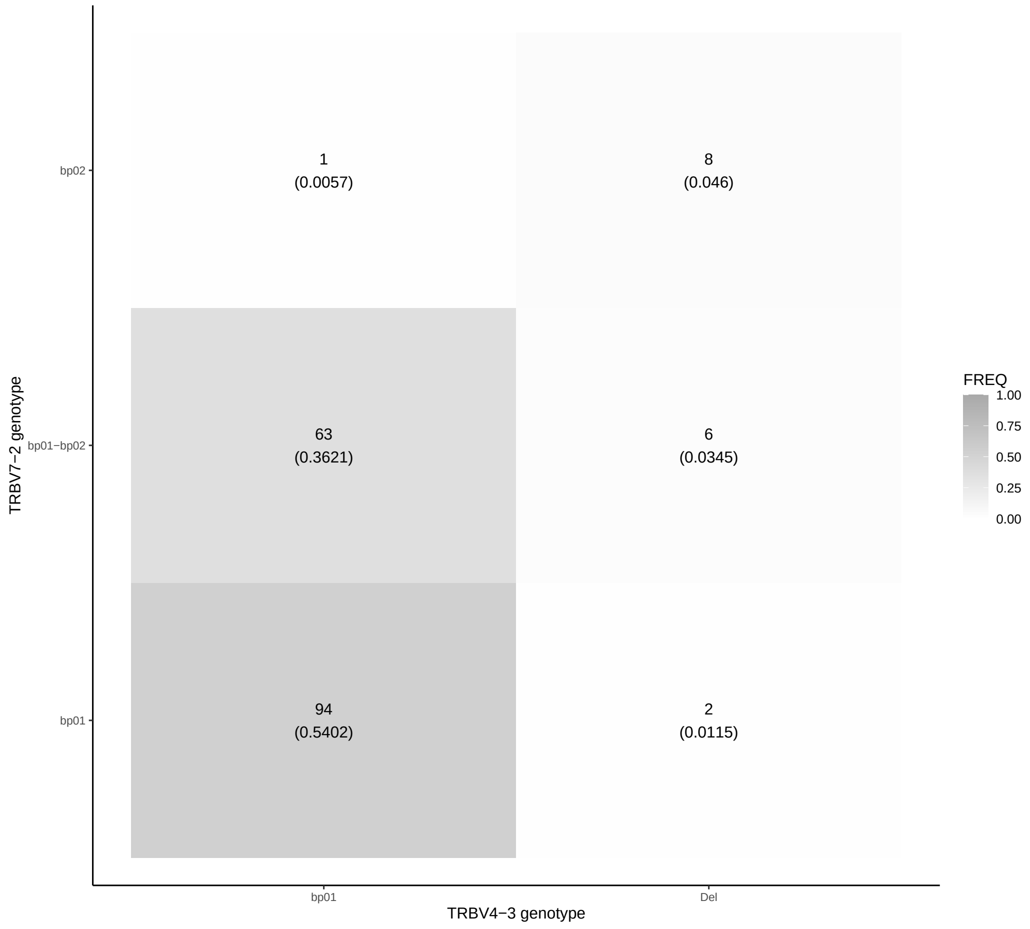


Fig S10: **Heatmap of TRBV4-3 genotype correlates to the TRBV7-2 genotype in DS3.** The X-axis represents the TRBV7-2 genotypes, the Y-axis is the TRBV4-3 genotypes, the values in each block are the number of individuals with the same inferred genotypes, and in brackets is the frequency of individuals who appear in this comparison.

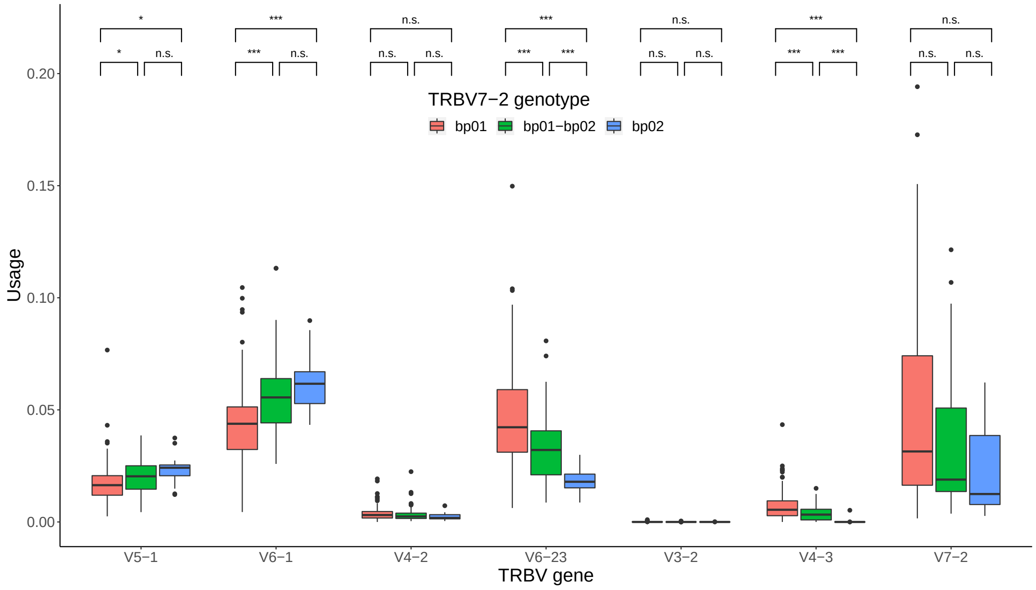


Fig S11: **TRBV gene usage according to the TRBV7-2 genotype in DS3.** Colors correspond to the TRBV7-2 genotype group. Statistical significance was determined using a Mann-Whitney test and adjusted by Bonferroni correction (see Methods; n.s. - not significant, * - $p < 0.05$, ** - $p < 0.01$, and *** - $p < 0.001$).

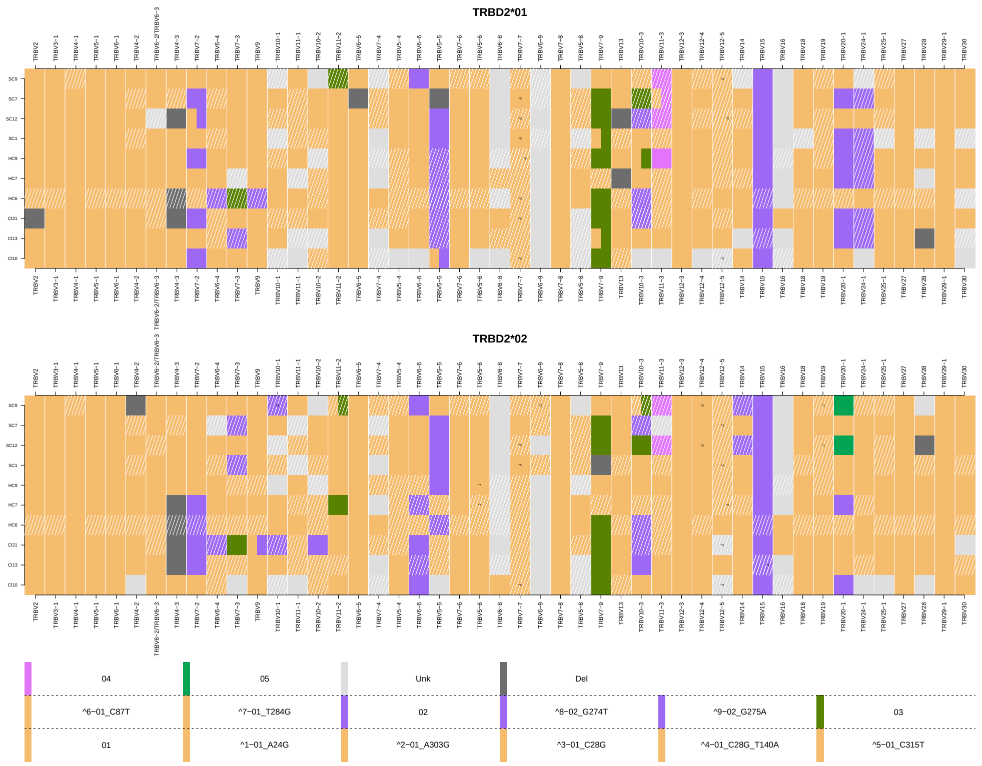


Fig S12: **TRBV haplotypes for 10 individuals from DS1.** The upper and lower panels show the TRBV haplotypes anchored with TRBD2*01 and TRBD2*02, respectively. Each row is an individual's haplotype, and each column is a V gene call. The colors correspond to the V alleles and the tile annotations correspond to the undocumented allele variations.

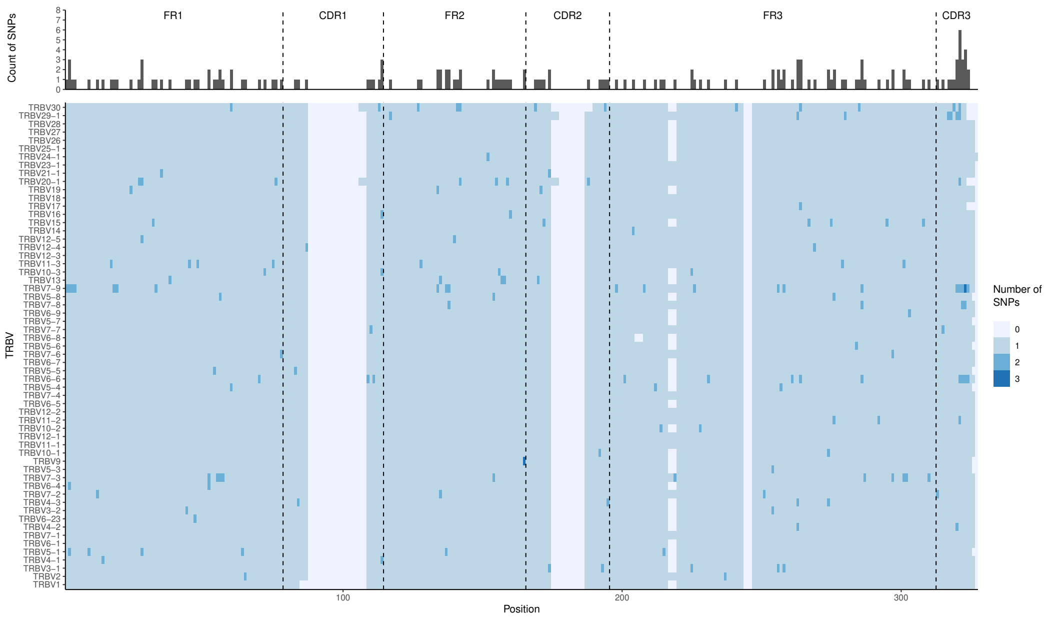


Fig S13: **TRBV SNPs distribution.** The top panel show the SNPs distribution across the V regions (FRX, CDRX) and position. The Y axis is the sum of the SNPs per the position in the X axis. The bottom panel show the distribution of the SNPs per V gene. The Y axis is the different V genes, the X axis is the positions across the V. The color scale represents the number of SNPs observed .

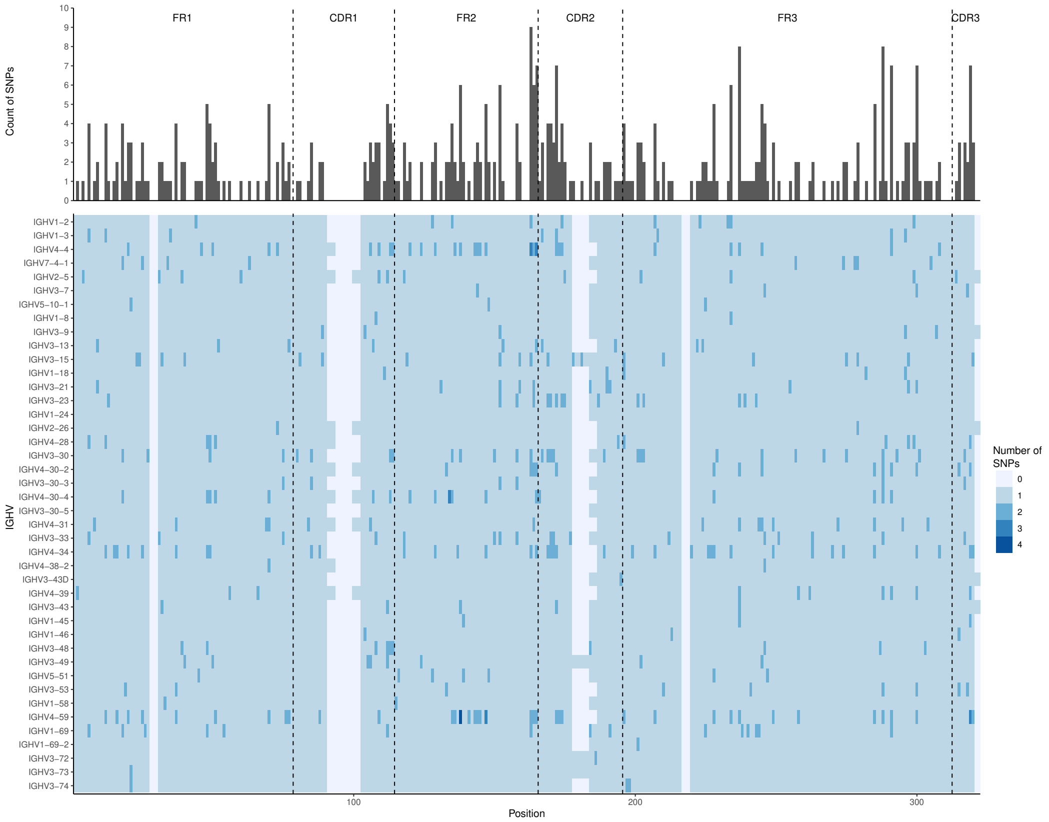


Fig S14: **IGHV SNPs distribution.** The top panel show the SNPs distribution across the V regions (FRX, CDRX) and position. The Y axis is the sum of the SNPs per the position in the X axis. The bottom panel show the distribution of the SNPs per V gene. The Y axis is the different V genes, the X axis is the positions across the V. The color scale represents the number of SNPs observed .

Table S1: **DS2 data-set sources citation.** The "Individuals" column displays the names we gave the individuals int the source. The "Citation" column is the citation of the paper the that data was obtained from and for the samples that were obtained from 10x Genomics is the citation that corresponds to the 10x Genomics requirements.

| Source | Individuals | Citations |
| --- | --- | --- |
| 10x Genomics [1] | sc5p | Human T cells from a Healthy Donor 1k cells (v2), Single Cell Immune Profiling Dataset by Cell Ranger 5.0.0, 10x Genomics, (2020, November 19). |
| | Parent_SC5 | PBMCs from a Healthy Donor: Paired T Cell Receptor, Single Cell Immune Profiling Dataset by Cell Ranger 4.0.0, 10x Genomics, (2020, July 7). |
| | donor1, donor2, donor3, and donor4 | CD8+ T cells of Healthy Donor 1-4, Single Cell Immune Profiling Dataset by Cell Ranger 3.0.2, 10x Genomics, (2019, May 9). |
| | pbmc2 | PBMCs of a healthy donor - TCR enrichment from amplified cDNA, Single Cell Immune Profiling Dataset by Cell Ranger 3.0.0, 10x Genomics, (2018, November 19). |
| | nsclc | NSCLC tumor - TCR enrichment from amplified cDNA, Single Cell Immune Profiling Dataset by Cell Ranger 3.0.0, 10x Genomics, (2018, August 1). |
| GEO [3] | C141, C142, and C143 | Liao et al. [8] |
| EGA [6] | ERS1, ERS2, ERS3, ERS4, ERS5, Healthy_Control_1, Healthy_Control_2, Healthy_Control_3, Healthy_Control_4, Healthy_Control_5, LRS2, LRS3, LRS4, and LRS5 | Wen et al. [11] |

Table S2: **BIOMED-2 allele patterns.**

| GENE | PATTERN | ALLELES |
|---|---|---|
| TRBV1 | bp01 | 01 |
| TRBV10-1 | bp01 | 01 |
| TRBV10-1 | bp02 | 02 |
| TRBV10-2 | bp01 | 01 |
| TRBV10-2 | bp02 | 02 |
| TRBV10-3 | bp01 | 01, 02, 04 |
| TRBV10-3 | bp02 | 03 |
| TRBV11-1 | bp01 | 01 |
| TRBV11-2 | bp01 | 01 |
| TRBV11-2 | bp02 | 02 |
| TRBV11-2 | bp03 | 03 |
| TRBV11-3 | bp01 | 01, 02, 04 |
| TRBV11-3 | bp02 | 03 |
| TRBV12-3/TRBV12-4 | bp01 | TRBV12-4*01, TRBV12-3*01 |
| TRBV12-3/TRBV12-4 | bp02 | TRBV12-4*02 |
| TRBV12-5 | bp01 | 01 |
| TRBV13 | bp01 | 01 |
| TRBV13 | bp02 | 02 |
| TRBV14 | bp01 | 01 |
| TRBV14 | bp02 | 02 |
| TRBV15 | bp01 | 01 |
| TRBV15 | bp02 | 02 |
| TRBV15 | bp03 | 03 |
| TRBV16 | bp01 | 01, 02 |
| TRBV16 | bp02 | 03 |
| TRBV18 | bp01 | 01 |
| TRBV19 | bp01 | 01 |
| TRBV19 | bp02 | 02, 03 |
| TRBV2 | bp01 | 01, 02 |
| TRBV2 | bp02 | 03 |
| TRBV20-1 | bp01 | 01, 02 |
| TRBV20-1 | bp02 | 03 |
| TRBV20-1 | bp03 | 04, 06 |
| TRBV20-1 | bp04 | 05 |

| | | |
|---|---|---|
| TRBV20-1 | bp05 | 07 |
| TRBV20/OR9-2 | bp01 | 01 |
| TRBV20/OR9-2 | bp02 | 02 |
| TRBV20/OR9-2 | bp03 | 03 |
| TRBV21-1 | bp01 | 01 |
| TRBV21-1 | bp02 | 02 |
| TRBV21/OR9-2 | bp01 | 01 |
| TRBV23-1 | bp01 | 01 |
| TRBV23/OR9-2 | bp01 | 01 |
| TRBV23/OR9-2 | bp02 | 02 |
| TRBV24-1 | bp01 | 01 |
| TRBV24-1 | bp02 | 02 |
| TRBV24/OR9-2 | bp01 | 01 |
| TRBV25-1 | bp01 | 01 |
| TRBV26 | bp01 | 01 |
| TRBV26/OR9-2 | bp01 | 01, 02 |
| TRBV27 | bp01 | 01 |
| TRBV28 | bp01 | 01 |
| TRBV29-1 | bp01 | 01 |
| TRBV29-1 | bp02 | 02 |
| TRBV29-1 | bp03 | 03 |
| TRBV29/OR9-2 | bp01 | 01 |
| TRBV29/OR9-2 | bp02 | 02 |
| TRBV3-1 | bp01 | 01 |
| TRBV3-1 | bp02 | 02 |
| TRBV3-2 | bp01 | 01 |
| TRBV3-2 | bp02 | 02, 03 |
| TRBV30 | bp01 | 01 |
| TRBV30 | bp02 | 02 |
| TRBV30 | bp03 | 03 |
| TRBV30 | bp04 | 04 |
| TRBV30 | bp05 | 05 |
| TRBV4-1 | bp01 | 01, 02 |
| TRBV4-2 | bp01 | 01 |
| TRBV4-2 | bp02 | 02 |
| TRBV4-3 | bp01 | 01, 04 |

| | | |
|---|---|---|
| TRBV4-3 | bp02 | 02 |
| TRBV4-3 | bp03 | 03 |
| TRBV5-1 | bp01 | 01 |
| TRBV5-1 | bp02 | 02 |
| TRBV5-3 | bp01 | 01 |
| TRBV5-3 | bp02 | 02 |
| TRBV5-4 | bp01 | 01, 03 |
| TRBV5-4 | bp02 | 02 |
| TRBV5-4 | bp03 | 04 |
| TRBV5-5 | bp01 | 01, 02, 03 |
| TRBV5-6 | bp01 | 01 |
| TRBV5-7 | bp01 | 01 |
| TRBV5-8 | bp01 | 01 |
| TRBV5-8 | bp02 | 02 |
| TRBV6-1 | bp01 | 01 |
| TRBV6-2/TRBV6-3 | bp01 | TRBV6-2*01, TRBV6-3*01 |
| TRBV6-4 | bp01 | 01, 02 |
| TRBV6-5 | bp01 | 01 |
| TRBV6-6 | bp01 | 01, 03 |
| TRBV6-6 | bp02 | 02 |
| TRBV6-6 | bp03 | 04 |
| TRBV6-6 | bp04 | 05 |
| TRBV6-7 | bp01 | 01 |
| TRBV6-8 | bp01 | 01 |
| TRBV6-9 | bp01 | 01 |
| TRBV7-1 | bp01 | 01 |
| TRBV7-2 | bp01 | 01, 04 |
| TRBV7-2 | bp02 | 02 |
| TRBV7-2 | bp03 | 03 |
| TRBV7-3 | bp01 | 01 |
| TRBV7-3 | bp02 | 02 |
| TRBV7-3 | bp03 | 03 |
| TRBV7-3 | bp04 | 04 |
| TRBV7-3 | bp05 | 05 |
| TRBV7-4 | bp01 | 01 |
| TRBV7-6 | bp01 | 01, 02 |

| | | |
|---|---|---|
| TRBV7-7 | bp01 | 01, 02 |
| TRBV7-8 | bp01 | 01 |
| TRBV7-8 | bp02 | 02 |
| TRBV7-8 | bp03 | 03 |
| TRBV7-9 | bp01 | 01, 02, 03 |
| TRBV7-9 | bp02 | 04 |
| TRBV7-9 | bp03 | 05 |
| TRBV7-9 | bp04 | 06 |
| TRBV7-9 | bp05 | 07 |
| TRBV9 | bp01 | 01 |
| TRBV9 | bp02 | 02 |
| TRBV9 | bp03 | 03 |

Table S3: **Adaptive allele patterns.**

| GENE | PATTERN | ALLELES |
|---|---|---|
| TRBV10-1 | ap01 | 01, 02 |
| TRBV10-2 | ap01 | 01, 02 |
| TRBV10-3 | ap01 | 01, 02, 03, 04 |
| TRBV11-1 | ap01 | 01 |
| TRBV11-2 | ap01 | 01, 03 |
| TRBV11-2 | ap02 | 02 |
| TRBV11-3 | ap01 | 01, 02, 04 |
| TRBV11-3 | ap02 | 03 |
| TRBV12-1 | ap01 | 01 |
| TRBV12-2 | ap01 | 01 |
| TRBV12-3/TRBV12-4 | ap01 | TRBV12-4*01, TRBV12-4*02, TRBV12-3*01 |
| TRBV12-5 | ap01 | 01 |
| TRBV13 | ap01 | 01, 02 |
| TRBV14 | ap01 | 01, 02 |
| TRBV15 | ap01 | 01 |
| TRBV15 | ap02 | 02 |
| TRBV15 | ap03 | 03 |
| TRBV16 | ap01 | 01, 02, 03 |
| TRBV17 | ap01 | 01, 02 |
| TRBV18 | ap01 | 01 |
| TRBV19 | ap01 | 01, 02, 03 |
| TRBV2 | ap01 | 01, 02, 03 |
| TRBV20-1 | ap01 | 01, 02, 03, 05, 06, 07 |
| TRBV20-1 | ap02 | 04 |
| TRBV20/OR9-2 | ap01 | 01, 02, 03 |
| TRBV21-1 | ap01 | 01, 02 |
| TRBV21/OR9-2 | ap01 | 01 |
| TRBV23-1 | ap01 | 01 |
| TRBV23/OR9-2 | ap01 | 01, 02 |
| TRBV24-1 | ap01 | 01, 02 |
| TRBV25-1 | ap01 | 01 |
| TRBV26 | ap01 | 01 |
| TRBV26/OR9-2 | ap01 | 01, 02 |

| | | |
|---|---|---|
| TRBV27 | ap01 | 01 |
| TRBV28 | ap01 | 01 |
| TRBV29-1 | ap01 | 01, 02 |
| TRBV29-1 | ap02 | 03 |
| TRBV29/OR9-2 | ap01 | 01, 02 |
| TRBV3-1/TRBV3-2 | ap01 | TRBV3-1*01, TRBV3-1*02, TRBV3-2*01, TRBV3-2*02, TRBV3-2*03 |
| TRBV30 | ap01 | 01, 03 |
| TRBV30 | ap02 | 02, 04 |
| TRBV30 | ap03 | 05 |
| TRBV4-1 | ap01 | 01, 02 |
| TRBV4-2 | ap01 | 01 |
| TRBV4-2 | ap02 | 02 |
| TRBV4-3 | ap01 | 01, 02, 03, 04 |
| TRBV5-1 | ap01 | 01, 02 |
| TRBV5-3 | ap01 | 01, 02 |
| TRBV5-4 | ap01 | 01, 02, 03, 04 |
| TRBV5-5 | ap01 | 01, 02, 03 |
| TRBV5-6 | ap01 | 01 |
| TRBV5-7 | ap01 | 01 |
| TRBV5-8 | ap01 | 01, 02 |
| TRBV6-1 | ap01 | 01 |
| TRBV6-2/TRBV6-3 | ap01 | TRBV6-2*01, TRBV6-3*01 |
| TRBV6-4 | ap01 | 01, 02 |
| TRBV6-5/TRBV6-6 | ap01 | TRBV6-5*01, TRBV6-6*01, TRBV6-6*02, TRBV6-6*03 |
| TRBV6-5/TRBV6-6 | ap02 | TRBV6-6*04 |
| TRBV6-5/TRBV6-6 | ap03 | TRBV6-6*05 |
| TRBV6-7 | ap01 | 01 |
| TRBV6-8 | ap01 | 01 |
| TRBV6-9 | ap01 | 01 |
| TRBV7-1 | ap01 | 01 |
| TRBV7-2 | ap01 | 01, 02, 04 |
| TRBV7-2 | ap02 | 03 |
| TRBV7-3 | ap01 | 01, 05 |

| | | |
|---|---|---|
| TRBV7-3 | ap02 | 02 |
| TRBV7-3 | ap03 | 03 |
| TRBV7-3 | ap04 | 04 |
| TRBV7-4 | ap01 | 01 |
| TRBV7-6 | ap01 | 01, 02 |
| TRBV7-7 | ap01 | 01, 02 |
| TRBV7-8 | ap01 | 01 |
| TRBV7-8 | ap02 | 02 |
| TRBV7-8 | ap03 | 03 |
| TRBV7-9 | ap01 | 01, 02, 03, 04 |
| TRBV7-9 | ap02 | 05 |
| TRBV7-9 | ap03 | 06 |
| TRBV7-9 | ap04 | 07 |
| TRBV9 | ap01 | 01, 02, 03 |

Table S4: The distribution parameters of the TRBD2*01 fraction according to the TRBD2 genotype group in DS4.

| TRBD2 genotype group \ TRBD2*01 fraction | Average | Median | Standard deviation |
|---|---|---|---|
| TRBD2*01 homozygous | 0.96 | 0.96 | 0.003 |
| TRBD2 heterozygous | 0.453 | 0.452 | 0.022 |
| TRBD2*02 homozygous | 0.127 | 0.127 | 0.008 |

Table S5: **Previously unknown alleles comparison.** The left column is the unknown alleles of DS1 and DS2, columns 2-5 indicate if the allele was found in another dataset, i.e., DS3, DS4, Lou et al. [9], and pmTRIG [5], respectively. Column 6 indicates if the undocumented allele was observed in the long-read assemblies. The colors red, green, and blue in the first column correspond to alleles not following the expected multi-modal distribution, alleles adjacent to nucleotide stretches, or both, respectively. The purple color in the first column corresponds to an allele found in more than one data-set. For DS3 and DS4, it is impossible to identify undocumented SNPs that are outside the amplified region. Hence, out of the 24 undocumented alleles, 7 and 18 were out of range for DS3 and DS4, respectively.

| DS1 [4] and DS2 [1] | DS3 [10] | DS4 [2] | Luo et al. [9] | pmTRIG [5] | long-read assemblies |
|---|---|---|---|---|---|
| TRBV10-1*02_G274T | TRBV10-1*bp02_G274T | | TRBV10-1*02_gt234E_ | | True |
| TRBV6-6*01_G264A | | | | | |
| TRBV7-6*01_G297A | | | | | |
| TRBV12-4*01_C87T | | | | TRBV12-4_3 | True |
| TRBV12-5*01_C28G | | | TRBV12-5*01_cg27HD | | True |
| TRBV12-5*01_C28G_T140A | | | | | |
| TRBV13*01_A170T | | | | | |
| TRBV13*01_T158C | | | | | |
| TRBV15*02_G275A | | | | | |
| TRBV19*01_A24G | | | TRBV19*01_ag23PP | TRBV19_3 | True |
| TRBV20-1*01_C142A | | | | TRBV20-1_4 | |
| TRBV30*01_A113C | | | | | |
| TRBV5-6*01_T284G | TRBV5-6*bp01_T284G | | TRBV5-6*01_tg244LW | TRBV5-6_4 | True |
| TRBV5-8*01_C56T_C276T | TRBV5-8*bp01_C276T | | TRBV5-8*01_ct55AV_ct236NN | | |
| TRBV6-2/TRBV6-3*01_G47A | | | | | |
| TRBV6-6*01_C261T | TRBV6-6*bp01_C261T | | | | |
| TRBV6-9*01_A303G | | | TRBV6-9*01_ag263VV | | |
| TRBV7-7*01_C315T | TRBV7-7*bp01_C315T | TRBV7-7*ap01_C315T | | TRBV7-7_2 | True |
| TRBV10-2*01_C214T | | | | TRBV10-2_3 | |
| TRBV29-1*01_A280C | | | TRBV29-1*01_ac246ML | TRBV29-1_2 | True |
| TRBV10-3*01_C225G | | | | | |
| TRBV4-1*01_C14T | | | | TRBV4-1_6 | |
| TRBV4-3*01_C274T | | | | | |
| TRBV7-3*01_G301A | | TRBV7-3*ap01_G301A | | TRBV7-3_5 | True |

Table S6: **Incomplete allele extensions table.** The left column is the call of the undocumented allele candidate, the second column is the matched incomplete allele reference in IMGT [7], and the third column is the length difference between the sequence of the undocumented allele candidate and the incomplete allele reference

| Undocumented allele candidate | Incomplete allele | Length difference |
|---|---|---|
| TRBV10-3*01_T114C_G156A | TRBV10-3*03 | 14 |
| TRBV14*01_G204A | TRBV14*02 | 5 |
| TRBV2*01_G65A | TRBV2*02 | 5 |
| TRBV20-1*05_A142C | TRBV20-1*02 | 5 |
| TRBV5-5*01_A54C | TRBV5-5*02 | 4 |

Table S7: **5′ UTR variants.** The left column is the allele for which the 5′ UTR variant was found. The second column indicates if more than one consensus sequence was found for the allele. The third column is the number of samples in the cluster. The fourth and fifth columns indicate the 5′UTR SNP position and change for L-PART2 and L-PART1. The sixth column indicates if the variation is from alternative splicing events and shows the alternative sequence.

| Allele | Consensus allele | cluster count | L-PART2 variation | L-PART1 variation | Alternative splicing variation |
|---|---|---|---|---|---|
| TRBV12-4*01 | TRBV12-4*01_2 | 22 | | G5A | |
| TRBV13*01 | TRBV13*01_2 | 9 | T1G | | |
| TRBV13*01 | TRBV13*01_3 | 25 | T1G | A53G | |
| TRBV18*01 | TRBV18*01_2 | 15 | | G13C | |
| TRBV19*01 | TRBV19*01_2 | 14 | | T37C | |
| TRBV19*01_A24G | | 6 | | T37C | |
| TRBV23-1*01 | TRBV23-1*01_1 | 17 | | | GACATTCTCTTTCTTTGTCTACGACATCTTTC TCAGGTCCTTTCTCGAATTGTTTGTTTTGTTT TGTTTTGTTTTGTTTTGTTTTGTTTCGAACCT AGACGACCAACCGTTCGGGTCCTGTTCCT AAAAAGAAAACACGAGTCCCGCGTA |
| TRBV23-1*01 | TRBV23-1*01_2 | 6 | | | GACATTCTCTTTCTTTGTCTACGACATCTTT CTCAGGTCCTTTCTCGAATTGTTTGTTTTGT TTTGTTTTGTTTTGTTTTGTTTTGTTTTGTT TCGAACCTAGACGACCAACCGTTCGGGTCCT GTTCCTAAAAAGAAAACACGAGTCCCGCGTA |
| TRBV25-1*01 | TRBV25-1*01_2 | 15 | | A25G | |
| TRBV28*01 | | 28 | | G18C | |
| TRBV5-4*01 | TRBV5-4*01_2 | 8 | | T26C | |
| TRBV5-8*01_C56T_C276T | TRBV5-8*01_C56T_C276T_2 | 2 | | C47T | |
| TRBV6-2/TRBV6-3*01 | TRBV6-2/TRBV6-3*01_2 | 10 | | G26A,G27C | |
| TRBV6-2/TRBV6-3*01 | TRBV6-2/TRBV6-3*01_3 | 8 | | C29T | |
| TRBV6-2/TRBV6-3*01_G47A | | 1 | | G26A,G27C | |
| TRBV6-9*01 | | 2 | | A47C,G48A | |
| TRBV6-9*01_A303G | | 2 | | A47C,G48A | |
| TRBV7-3*02 | | 4 | | G12C | |
| TRBV7-7*01 | | 18 | | | ATTCTGTTTCCACAG |
| TRBV7-7*01_C315T | TRBV7-7*01_C315T_1 | 16 | | | ATTCTGTTTCCACAG |

Table S8: **5′ UTR undocumented sequences.** The left column is the allele for which the 5′ UTR variant was undocumented. The second column indicates which L-PART sequence is absent from IMGT. The third column is the number of samples in the cluster. The fourth column indicates which L-PART is also observed in the long read genomic assembly data-set.

| Allele | Absent sequences | cluster count | Observed sequences in long-read assemblies |
|---|---|---|---|
| TRBV10-2*02 | L-PART1 and L-PART2 | 7 | L-PART1 and L-PART2 |
| TRBV10-3*02 | L-PART1 | 19 | |
| TRBV10-3*03 | L-PART1 and L-PART2 | 11 | L-PART2 |
| TRBV11-3*04 | L-PART1 and L-PART2 | 11 | L-PART1 and L-PART2 |
| TRBV14*02 | L-PART1 and L-PART2 | 8 | L-PART2 |
| TRBV15*02 | L-PART1 and L-PART2 | 1 | L-PART2 |
| TRBV15*02_G275A | L-PART1 and L-PART2 | 1 | |
| TRBV2*02 | L-PART1 and L-PART2 | 1 | L-PART1 and L-PART2 |
| TRBV20-1*05 | L-PART1 and L-PART2 | 6 | L-PART1 |
| TRBV3-2*03 | L-PART1 and L-PART2 | 2 | |
| TRBV5-5*02 | L-PART1 and L-PART2 | 23 | |

Table S9: **Undocumented allele verification in artificial partial libraries.** To assess the ability of the presented pipeline to infer partial TRBV undocumented alleles from partial VDJ sequences, we generated from DS1 two artificial data-sets that simulate TRB repertoires obtained by BIOMED-2 and Adaptive Biotechnologies. For generating DS1 to stimulate a data-set obtained by BIOMED-2 protocol sequencing, the input sequences after the first alignment (see methods section 3.4) were trimmed by using the TRBV germline positions that the BIOMED-2 primers bind to. Sequences that aligned to TRBV genes whose BIOMED-2 primers do not bind were filtered out. Generating DS1 as a data-set obtained by Adaptive Biotechnologies pipeline was done by taking 87 nt up to the 23rd germline position from the end of the TRBJ gene whose sequence was assigned to. Next, we ran the trimmed sequences of both simulated data-sets through the pipeline. The table show the successful identification of the full length alleles by the artificial partial libraries. The left column is the undocumented allele from the complete DS1, the second and third columns are the artificial DS1 undocumented alleles for BIOMED2 and Adaptive lengths, respectively. The † indicated an undocumented allele inference which is identical to a known allele within the reference and hence did not make it to the genotype. The purple color indicates an allele that carries a SNP that is also found within the full length sequence. However, due to noise in the beginning of the full length sequences, there is no exact match to the undocumented allele version. This prevents the undocumented allele from passing the filtration step and enter the genotype, the same as in the artificial BIOMED2 dataset. This issue is the outcome of the initial sequence length and will persist in non-artificial sequences.

| DS1 [4] undocumented alleles | DS1 [4] as BIOMED2 | DS1 [4] as Adaptive |
|---|---|---|
| TRBV12-5*01_C28G | not applicable | not applicable |
| TRBV5-8*01_C56T_C276T | TRBV5-8*bp01_C276T | not applicable |
| TRBV6-9*01_A303G | TRBV6-9*bp01_A303G | TRBV6-9*ap01_A303G† |
| TRBV7-7*01_C315T | TRBV7-7*bp01_C315T | TRBV7-7*ap01_C315T |
| TRBV12-4*01_C87T | not applicable | not applicable |
| TRBV19*01_A24G | not applicable | not applicable |
| TRBV6-2/TRBV6-3*01_G47A | not applicable | not applicable |
| TRBV15*02_G275A | TRBV15*bp02_G275A | not applicable |
| TRBV12-5*01_C28G_T140A | not applicable | not applicable |
| TRBV5-6*01_T284G | TRBV5-6*bp01_T284G | not applicable |
| TRBV10-1*02_G274T | TRBV10-1*bp02_G274T | not applicable |
| TRBV6-6*01_G264A | TRBV6-6*bp01_G264A | not applicable |
| | TRBV10-2*bp01_T233C | |
| | TRBV6-9*bp01_A229C_A303G | |

Table S10: **Previously unknown alleles comparison.** The left column is the unknown alleles of DS3, columns 2-5 indicate if the allele was found in another dataset, DS1 and DS2, DS4, Lou et al. [9], and pmTRIG [5], respectively. Column 6 indicates if the undocumented allele was observed in long-read assemblies. The colors red, green, and blue in the first column correspond to alleles not following the expected multi-modal distribution, alleles adjacent nucleotide stretches, or both, respectively. The purple color in the first column corresponds to an allele found in more than one dataset.

| DS3 [10] | DS1 [4] and DS2 [1] | DS4 [2] | Luo et al. [9] | pmTRIG [5] |
|---|---|---|---|---|
| TRBV10-1*bp01_C190A_C195T_A199G | | | | |
| TRBV10-1*bp01_G274T | | | | |
| TRBV10-1*bp02_G156A_G274T | | | | |
| TRBV10-1*bp02_G274T | TRBV10-1*02_G274T | | TRBV10-1*02_gt234 | |
| TRBV11-1*bp01_C149T | | | | TRBV11-1_5 |
| TRBV11-3*bp01_G297A | | | | |
| TRBV15*bp02_G153T | | | | |
| TRBV19*bp01_T310C_G311C_C314T | | | | |
| TRBV24-1*bp01_A316C | | | | |
| TRBV24-1*bp01_G252A | | | | |
| TRBV29-1*bp03_C315T | | | | |
| TRBV30*bp01_C253T | | | | |
| TRBV30*bp01_G298A | | TRBV30*ap01_G298A | | TRBV30_8 |
| TRBV30*bp02_C237T | | | | |
| TRBV5-4*bp01_G205A | | | | |
| TRBV5-4*bp01_G205A | | | | |
| TRBV5-5*bp01_T284G_G303C | | | | |
| TRBV5-6*bp01_T284G | TRBV5-6*01_T284G | | TRBV5-6*01_tg244 | TRBV5-6_4 |
| TRBV5-8*bp01_C276T | TRBV5-8*01_C56T_C276T | | TRBV5-8*01_ct236 | |
| TRBV6-4*bp01_C269G | | | | |
| TRBV6-5*bp01_G279A | | | | |
| TRBV6-6*bp01_C261T | TRBV6-6*01_C261T | | | TRBV6-6_8 |
| TRBV6-6*bp01_G256T | | | | TRBV6-6_4 |
| TRBV6-9*bp01_G155T_C156G_A303G | | | | |
| TRBV7-4*bp01_G251A | | | TRBV7-4*01_ga214 | TRBV7-4_3 |
| TRBV7-4*bp01_T306C_C307T | | | | |
| TRBV7-7*bp01_C315T | TRBV7-7*01_C315T | TRBV7-7*ap01_C315T | | TRBV7-7_2 |
| TRBV7-7*bp01_T273C | | | | |
| TRBV7-9*bp04_T312A | | | | |

Table S11: **Previously unknown alleles comparison.** The left column is the unknown alleles of datasets DS4, columns 2-5 indicates if the allele was found in another dataset, DS1 and DS2, DS3, Lou et al. [9], and pmTRIG [5], respectively. Column 6 indicates if the unknown allele was observed in long-read assemblies. The colors red, green, and blue in the first column correspond to alleles not following the expected multi-modal distribution, alleles adjacent nucleotide stretches, or both, respectively. The purple color in the first column corresponds to an allele found in more than one dataset.

| DS4 [2] | DS1 [4] and DS2 [1] | DS3 [10] | Luo et al. [9] | pmTRIG [5] | long-read assemblies |
|---|---|---|---|---|---|
| TRBV11-3*ap01_T312C | | | | | |
| TRBV15*ap02_C296T | | | | | |
| TRBV15*ap02_G290C | | | | | |
| TRBV20-1*ap01_G314A | | | | | |
| TRBV20-1*ap02_T310G | | | | | |
| TRBV25-1*ap01_A293G | | | | TRBV25-1_5 | |
| TRBV28*ap01_A297G | | | | | |
| TRBV30*ap01_G298A | | TRBV30*bp01_G298A | | TRBV30_8 | |
| TRBV4-3*ap01_A305C_T306C | | | | | |
| TRBV4-3*ap01_A305C_T306C_T308C | | | | | |
| TRBV4-3*ap01_G311C_G313C | | | | | |
| TRBV4-3*ap01_T308C_G311C | | | | | |
| TRBV4-3*ap01_T308C_T310C_G311C | | | | | |
| TRBV6-8*ap01_A293G | | | TRBV6-8*01_ag250 | | |
| TRBV7-3*ap01_G292T | | | TRBV7-3*01_gt255 | TRBV7-3_4 | True |
| TRBV7-3*ap01_G301A | TRBV7-3*01_G301A | | | TRBV7-3_5 | |
| TRBV7-4*ap01_G291C_A297G | | | | | |
| TRBV7-4*ap01_G291C_A297G_C314T | | | | | |
| TRBV7-6*ap01_C314G | | | | | |
| TRBV7-7*ap01_C307A_C315T | | | | | |
| TRBV7-7*ap01_C307T_C315T | | | | | |
| TRBV7-7*ap01_C315T | TRBV7-7*01_C315T | | | TRBV7-7_2 | |
| TRBV7-8*ap01_T295C | | | TRBV7-8*01_tc258 | | |
| TRBV7-9*ap01_G313T | | | | | |

[5] Indu Khatri, Magdalena A. Berkowska, Erik B. van den Akker, Cristina Teodosio, Marcel J.T. Reinders, and Jacques J.M. van Dongen. Population matched (pm) germline allelic variants of immunoglobulin (ig) loci: New pmig database to better understand ig repertoire and selection processes in disease and vaccination. *bioRxiv*, 2020.

[6] Ilkka Lappalainen, Jeff Almeida-King, Vasudev Kumanduri, Alexander Senf, John Dylan Spalding, Gary Saunders, Jag Kandasamy, Mario Caccamo, Rasko Leinonen, Brendan Vaughan, et al. The european genome-phenome archive of human data consented for biomedical research. *Nature genetics*, 47(7):692–695, 2015.

[7] Marie-Paule Lefranc, Véronique Giudicelli, Chantal Ginestoux, Julia Bodmer, Werner Müller, Ronald Bontrop, Marc Lemaitre, Ansar Malik, Valérie Barbié, and Denys Chaume. Imgt, the international immunogenetics database. *Nucleic Acids Research*, 27(1):209–212, 1999.

[8] Mingfeng Liao, Yang Liu, Jing Yuan, Yanling Wen, Gang Xu, Juanjuan Zhao, Lin Cheng, Jinxiu Li, Xin Wang, Fuxiang Wang, Lei Liu, Ido

Amit, Shuye Zhang, and Zheng Zhang. Single-cell landscape of bronchoalveolar immune cells in patients with covid-19. *Nature Medicine*, 26(6):842–844, Jun 2020.

[9] Shishi Luo, Jane A Yu, Heng Li, and Yun S Song. Worldwide genetic variation of the ighv and trbv immune receptor gene families in humans. *Life Science Alliance*, 2(2), 2019.

[10] Donjete Simnica, Nuray Akyüz, Simon Schliffke, Malte Mohme, Lisa v.Wenserski, Thorben Mährle, Lorenzo F. Fanchi, Katrin Lamszus, and Mascha Binder. T cell receptor next-generation sequencing reveals cancer-associated repertoire metrics and reconstitution after chemotherapy in patients with hematological and solid tumors. *OncoImmunology*, 8(11):e1644110, 2019. PMID: 31646093.

[11] Wen Wen, Wenru Su, Hao Tang, Wenqing Le, Xiaopeng Zhang, Yingfeng Zheng, Xiuxing Liu, Lihui Xie, Jianmin Li, Jinguo Ye, Liwei Dong, Xiuliang Cui, Yushan Miao, Depeng Wang, Jiantao Dong, Chuanle Xiao, Wei Chen, and Hongyang Wang. Immune cell profiling of covid-19 patients in the recovery stageby single-cell sequencing. *Cell Discovery*, 6(1):31, May 2020.
